# Development and validation of highly selective monoclonal antibodies for the detection of huntingtin neoepitopes

**DOI:** 10.64898/2026.08.21.746138

**Authors:** Antonino Missineo, Licia Tomei, Nadine Alaimo, Paola Martufi, Marta Zavattieri, Valeria Colicchia, Cristina Cariulo, Valentina Fodale, Jacqueline Séguin, Candi Esquina, Nian Huang, Hsuan-Yi Wu, James Pace, Jemima Phillips, Christian Landles, Celia Dominguez, Ignacio Munoz-Sanjuan, Elizabeth M. Doherty

## Abstract

Huntington’s disease is caused by a CAG repeat tract expansion in the huntingtin gene, resulting in production of pathogenic N-terminal huntingtin protein fragments associated with disease pathology. Despite their central role, detection of these fragments has relied on a limited antibody repertoire with reproducibility concerns. Here, we describe the generation and characterization of recombinant rabbit monoclonal antibodies targeting two reciprocal neoepitopes flanking the huntingtin exon 1/exon 2 junction corresponding to amino acids P90 and K91. The P90 antibodies (clones 1B12, 11G2) demonstrate fragment-length-selective recognition of the C-terminal HTTexon1 P90 neoepitope with no detectable binding to full length huntingtin. A side-by-side comparison of the widely used monoclonal antibody MW8 from two different sources revealed measurable lot-to-lot drift in its fragment selectivity, whereas the recombinant P90 antibodies, expressed from a defined, sequenced clone, maintained consistent specificity, addressing this source-dependent variability. Whereas P90-positive fragments can arise through alternative splicing of the *HTT1a* transcript, generation of the reciprocal K91 N-terminal HTTexon2 neoepitope would require site-specific proteolytic cleavage, a mechanism that has not yet been directly tested for lack of a suitable reagent. The K91 antibody (clone 7G10) binds the N-terminal K91 neoepitope with high affinity and specificity over full length huntingtin and provides, for the first time, a tool capable of directly interrogating whether such cleavage occurs. Neoepitope specificity of these antibodies was orthogonally confirmed by protease digestion (Lys-N and Arg-C) coupled with intact mass spectrometry. As an additional outcome of the immunization and selection strategy, we discovered human-mouse cross-reactive antibodies (clones 27F5, 31C10) targeting the proline-rich domain of huntingtin that will facilitate mouse-human translational studies. All antibodies are recombinant, ensuring long-term reproducibility, and are being made available, along with their sequences, to the research community.

## Introduction

Huntington’s disease (HD) is a devastating autosomal dominant neurodegenerative disorder characterized by progressive motor dysfunction, cognitive decline, and psychiatric disturbances [1]. The disease is caused by expansion of a CAG-repeat tract in exon 1 of the huntingtin (*HTT*) gene, which translates into an abnormally long polyglutamine (polyQ) tract in the full-length huntingtin protein (FL HTT) [2, 3]. Individuals with 40 or more CAG repeats invariably develop HD, with sign/symptom onset, severity, and progression inversely correlated with CAG-repeat length [1].

Accumulating evidence demonstrates that N-terminal fragments of huntingtin, particularly those derived from exon 1, corresponding to HTT aa 1-90 (HTTexon1), are highly pathogenic and play a critical role in HD molecular pathology [4]. The identification of such fragments has a rich history rooted in neuropathological and biochemical investigations. Early immunohistochemical studies of postmortem HD brain tissue identified huntingtin-immunoreactive aggregates in both neuronal nuclei and neuropil, suggesting that N-terminal HTT fragments were selectively sequestered into pathological inclusions in affected regions [5]. Biochemical analysis of human HD brain subsequently confirmed that enhanced proteolytic processing of mHTT occurs *in vivo*, with elevated levels of N-terminal HTT fragments detected in HD striatum relative to control tissue, pointing to the diseased brain environment as a driver of HTT cleavage [6]. Building on these observations, Lunkes and colleagues used cell-based models to characterize two short N-terminal cleavage products of mHTT, designated cp-A and cp-B, demonstrating that these proteolytically derived fragments differentially accumulated in cytoplasmic versus nuclear inclusions and that the smaller cp-A species preferentially targeted the nucleus [7]. Subsequent studies identified additional N-terminal fragments of similar apparent molecular weight, termed cp-1 and cp-2, in an inducible cell model expressing full-length mHTT; importantly, these fragments were generated through a caspase-independent proteolytic mechanism distinct from that producing cp-A and cp-B, and were detectable without pharmacological manipulation of the proteasome, suggesting they represent a physiologically relevant cleavage pathway [8]. For many years, N-terminal HTT fragments of the size of the HTTexon1 protein were assumed to arise exclusively through such proteolytic mechanisms; a pivotal discovery revealed, however, that *HTT* exon 1 does not always splice to exon 2, resulting in alternative processing that generates a short, polyadenylated mRNA transcript referred to as *HTT1a* [9]. This alternatively processed mRNA encodes the highly pathogenic protein fragment, HTT1a, comprising the N-terminal 17 amino acids, the expanded polyglutamine tract, and a proline-rich domain (PRD) that terminates in a C-terminal proline residue at amino acid position 90 (P90). The *HTT1a* transcript is produced in a CAG repeat length-dependent manner and has been detected in knock-in mouse models, patient-derived fibroblasts, and postmortem brain from individuals with both adult-onset and juvenile-onset HD [10–11].

Despite the central role in disease pathology attributed to HTTexon1, the field has historically relied on a limited repertoire of antibodies with significant limitations. Existing antibodies were typically designed to recognize specific domains, conformations, or aggregates [5, 12–15], rather than to define specific fragments via neoepitopes. In many cases, antibodies were chosen based solely on immunoblotting and were not subjected to rigorous validation prior to widespread use in alternative applications (e.g., immunoassays, immunoprecipitation). Furthermore, several commonly used HTTexon1 antibodies were derived from mouse hybridoma lines, which are susceptible to clonal drift and variability over time.

To address these limitations, we generated high-quality, well-characterized recombinant monoclonal antibodies (mAbs) specific for the HTTexon1 P90 C-terminal neoepitope as well as the complementary HTTexon2 K91 N-terminal neoepitope, corresponding to the reciprocal products of proteolytic cleavage at the P90–K91 site. We chose recombinant antibody production over hybridoma to ensure long-term consistency, reproducibility, and performance across applications. We describe here our immunization strategy, clone selection process, and characterization of these novel HTT mAbs by western blot (WB), surface plasmon resonance (SPR), and immunocytochemistry (ICC).

## Materials and Methods

### Recombinant Proteins

The following recombinant protein standards were produced and purified according to Pace *et al* [16]: FL Q23 HTT, human (CHDI-90001858); FL Q48 HTT, human (CHDI-90002137); FL Q7 HTT, mouse (CHDI-90002925). HTT Q23 Exon1 (CHDI-90001323); HTT Q43 Exon1 (CHDI-90001324); and fibrils formed from HTT Q43 Exon1 were produced according to the method of Reif *et al* [17]. Recombinant HTT fragment, aa 91-171, with a C-terminal maltose binding protein fusion (HTT 91-171 C-term MBP, CHDI-90004104) was produced by a method analogous to Reif *et al* [17]. Amino acid sequences for recombinant proteins are provided in the Supporting Information (S1).

### Plasmids

For over-expression studies, DNA was produced by de novo synthesis in a pcDNA3.1 mammalian expression vector (Invitrogen): Htt-Q23-pcDNA3.1, 1-90, human (CHDI-90003891); Htt-Q48-pcDNA3.1, 1-87, human (CHDI-90003881); Htt-Q48-pcDNA3.1, 1-89, human (CHDI-90003883); Htt-Q48-pcDNA3.1, 1-90, human (CHDI-90003884); Htt-Q48-pcDNA3.1, 1-91, human (CHDI-90003885); Htt-Q48-pcDNA3.1, 1-93, human (CHDI-90003887); Htt-Q48-pcDNA3.1, 1-95, human (CHDI-90003888); Htt-Q48-pcDNA3.1, 1-99, human (CHDI-90003890). Insert sequences with corresponding expressed protein sequences are provided in the Supporting Information (S2).

### Antibodies

The following antibodies were obtained from CHDI: 2B7 (CHDI-90000830); 4C9 (CHDI-90000833); MW8 (CHDI-90000942). Commercial antibodies MAB5490 and MABN2529 are available from Merck.

### Recombinant antibody production

Recombinant mAbs 1B12, 11G2, 7G10, 27F5, and 31C10 were produced at ABclonal (Woburn, MA). In brief, peptide antigens (Fig 1) were conjugated to keyhole limpet hemocyanin (KLH) for immunization; corresponding positive and negative counter-screening peptides were biotinylated (Fig 1) for ELISA-based screening. Antigen-directed B-cells were isolated by fluorescence activated cell sorting (FACS), cultured, and supernatants screened by ELISA at ABclonal. Selected clones were sequenced and provided as recombinant mAbs.

**Fig 1.**
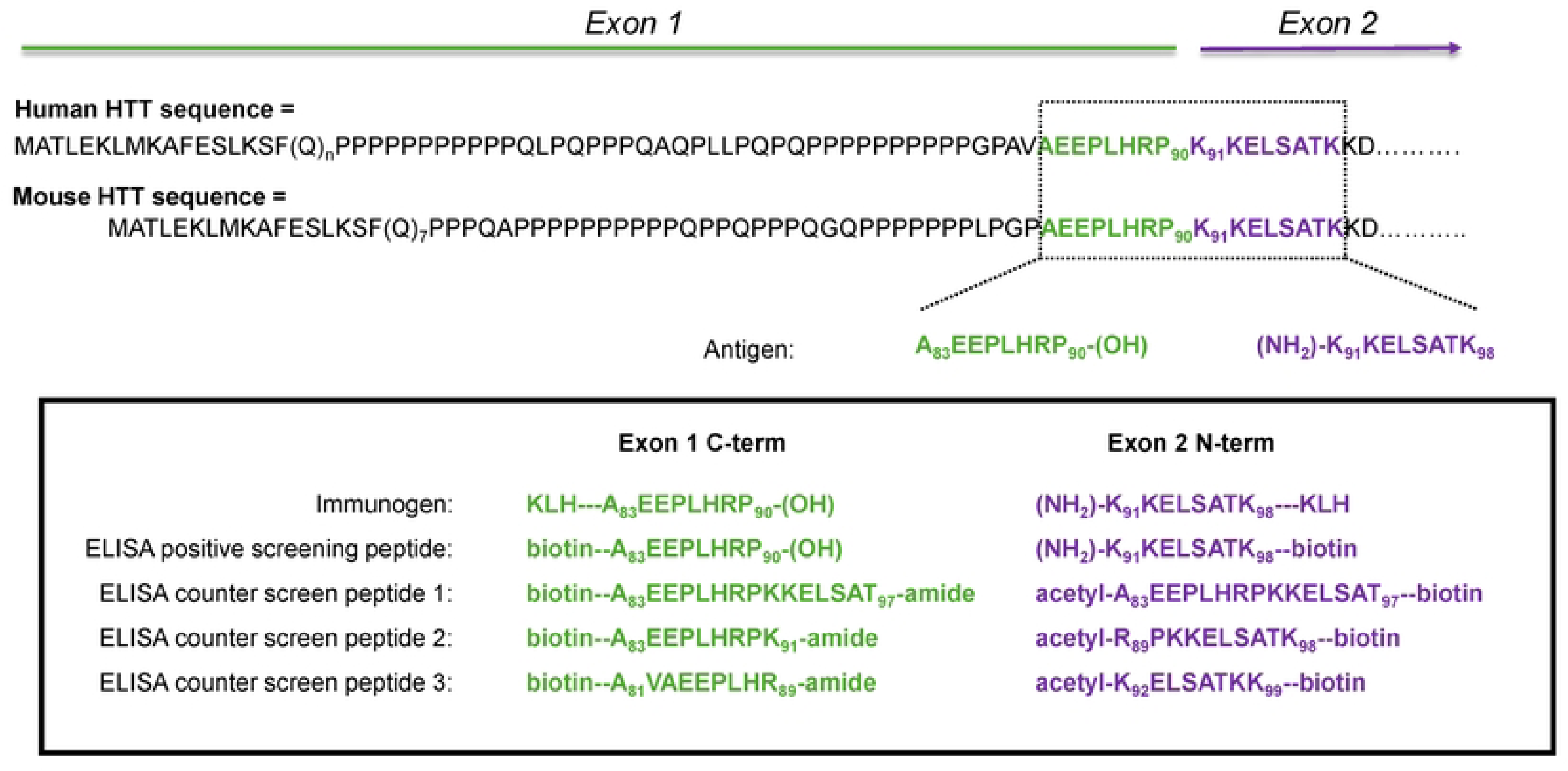
Human HTT N-terminal sequence compared to mouse HTT sequence (numbering based on Q23 for both sequences). Shared homologous sequence highlighted in dashed box. Peptide antigen sequences for HTTexon1 C-term (ending at P90) and HTTexon2 N-terminus (beginning at K91). Immunogen, positive and counter screen ELISA peptides design (in box). KLH = Keyhole Limpet Hemocyanin conjugate; (OH) = free carboxyl C-terminus; (NH_2_) = free amine N-terminus; acetyl = N-terminus acetylation; amide = C-terminus amide; dashed lines indicate linker.

### Western blot analysis of recombinant proteins

The recombinant proteins (FL Q23 HTT, human; HTT, 91-171, C-term MBP; HTT Q23 Exon1) were denatured at 95 °C in 4× Loading Buffer (125 mM Tris⋅HCl, pH 6.8, 6% SDS, 4 M urea, 4 mM EDTA, 30% glycerol, 4% 2-mercaptoethanol and bromophenol blue) and loaded on NuPAGE 4–12% Bis–Tris Gel (Thermo Fisher Scientific, catalog #WG1402BOX) at a concentration of 100 ng and separated by SDS-PAGE in MES-SDS running buffer at 120 V for 60 min. Proteins were transferred on PVDF membrane (Bio-Rad Laboratories, catalog #162–0177) using wet blotting (400mA for 90 minutes on ice). The membranes were blocked in 5% non-fat milk in Tris buffer saline pH 7.4 with 0.1% Tween-20 (TBS-T), for 1 h, at 25 °C, under gentle agitation. The mAbs 4C9, MW8, 1B12, 11G2, MAB5490 and 7G10 were prepared in 5% milk, TBS-T at a concentration of 1 μg/mL, then incubated with the membranes at 4 °C, for 16 h, under agitation. After several washes with TBS-T, the membranes were incubated for 45 min at 25 °C with either Goat Anti-Rabbit IgG Antibody, HRP-conjugate (cat # 12-348, Merck) or Goat Anti-mouse IgG antibody, HRP-conjugate (cat # A2554, Sigma Aldrich) prepared at dilution 1:5000 in 5% milk, TBS-T. Membranes were developed using Amersham™ ECL™ Western Blotting Detection Reagent (RPN2106, Cytvia) according to manufacturer’s instructions.

### Measurement of mAb affinity and binding kinetics to HTT recombinant proteins by surface plasmon resonance analysis

All surface plasmon resonance (SPR) experiments were performed using a Biacore T200 instrument (Cytvia), at 25 °C and data collected at 10 Hz. Rabbit mAbs 1B12, 11G2, 7G10, 27F5, 31G10 and MW8 were captured on Series S Sensor Chip Protein G (Cod. 29179315, Cytiva); mouse mAb 4C9 was captured on Series S Sensor Chip CM5 (Cod. BR100530, Cytiva) according to the Mouse Antibody Capture kit instructions (Cod. BR100838, Cytiva). For each experiment, one flow cell with only the capture molecule (protein G or anti-mouse IgGs) was left blank to serve as reference surface. Two buffer blanks were injected at the beginning and at the end of the analyte titration. Sensorgrams were double referenced, then data were fitted to a simple 1:1 interaction model using the local data analysis option of Biacore T200 Evaluation Software v. 3.2.1.

#### Binding to monomeric HTT Q23 Exon 1

The mAbs 1B12, 11G2, MW8 and 4C9 were prepared at a concentration of 10 μg mL^−1^ in 10 mM phosphate pH 7.4, 150 mM NaCl, 0.05% Tween-20 (PBS-P+) and injected for 60 s at 10 μL min^−1^ on the active flow cell. Recombinant HTT Q23 Exon 1 protein (lyophilized powder) was freshly disaggregated with trifluoracetic acid (TFA) and formulated at 20 µM in PBS pH 7.4. To evaluate binding to mAbs 1B12 and 11G2, the analyte solution in PBS-P+ was injected over the flow cells at concentrations from 100 nM to 3.12 nM (1:2 dilution, six-point titration). For mAbs MW8 and 4C9, an analyte concentration range of 5000 nM to 78 nM (1:2 dilution, seven-point titration) or 5000 nM to 6.9 nM (1:3 dilution, seven-point titration) was used, respectively. The complex was allowed to associate for 200 s and and dissociate for 600 s at a flow rate of 30 µL min^−1^. The surface was regenerated with two injections of 10 mM glycine pH 1.7 over 30 s.

#### Binding to fibrils

The HTT Q43 Exon1 fibrils were immobilized by amine coupling chemistry on Series S Sensor Chip CM5 (Cod. BR100530, Cytvia). The surfaces of two flow cells were activated for 7 min with a 1:1 mixture of 0.1 M N-hydroxysuccinimide and 0.1 M 3-(N,N-dimethylamino)propyl-N-ethylcarbodiimide at a flow rate of 5 μL min^−1^. The fibril solution was diluted 1:200 in sodium acetate pH 4.0 and immobilized at a density of 20 response units (RU) on flow cell 2; flow cell 1 was left blank to serve as reference surface. Both surfaces were blocked with a 7 min injection of 1 M ethanolamine, pH 8.0. To collect kinetic binding data the mAbs, in 10 mM HEPES, pH 7.4, 150 mM NaCl, 0,05% Tween-20 (HBS-P+), were injected over the active and reference flow cells at concentrations of 100 nM to 0.16 nM (1:5 dilution, five-point titration) at a flow rate of 30 μL min^−1^. The complex was allowed to associate for 200 s and and dissociate for 900 s at a flow rate of 30 µL min^−1^. The surfaces were regenerated with one injection of 10 mM glycine pH 1.7 over 30 s.

#### Binding to HTT, 91-171, C-term MBP

The mAb 7G10 was prepared at a concentration of 10 μg mL^−1^ in PBS-P+ and injected on the active flow cell for 60 s at 10 μL min^−1^. Recombinant HTT, 91-171, C-term MBP, was centrifuged at 17530 × *g*, at 4 °C for 5 min, then diluted in PBS-P+ and injected over the flow cells at concentrations from 100 nM to 3.12 nM (1:2 dilution, six-point titration). The complex was allowed to associate for 200 s and dissociate for 600 s at a flow rate of 30 µL min^−1^. The surface was regenerated with two injections of 10 mM glycine pH 1.7 over 30 s.

#### Binding to FL HTT

The mAbs 4C9, MW8, 1B12, 11G2, 7G10, 27F5, and 31C10 were prepared at 1 μg mL^−1^ in PBS-P+ and injected on active flow cells for 60 s at 10 μL min^−1^. Recombinant FL Q23 HTT (human) and FL Q7 HTT (mouse) was centrifuged at 17530 × *g*, at 4 °C for 5 min, then diluted in PBS-P+ and injected over the flow cells at concentrations from 300 nM to 1.2 nM (1:2 dilution, eight-point titration). The complex was allowed to associate for 200 s and dissociate for 900 s at a flow rate of 30 µL min^−1^. The surface was regenerated with two injections of 10 mM glycine pH 1.7 over 30 s.

### Epitope mapping by western blot and filter trap

#### Transfection Protocol

HEK293 cells were grown and maintained in Gibco Expi293 expression medium (cat # A14351-01, ThermoFisher) with 0.5X Gibco Penicillin-Streptomycin (P/S, cat # 15140-122, ThermoFisher). HEK293 cells (2 mL at 1.2 X 10^6^ cells/mL) in Expi293 medium without P/S were prepared in each well of a 24-well plate (cat # 931565-G-1X, Thomson Solutions at Work). The 24-well plate was covered with sterile AeraSeal^TM^ film (cat # A9224, Excel Scientific, Inc.) and kept at 37 °C, 5% CO_2_, under gentle agitation. The following day, a transfection mixture was prepared using DNA and FuGENE HD reagent (cat # E2311, Promega) at a DNA to reagent ratio of 1 µg to 2 µL (2 µg of plasmid to 6 µL of reagent) with a final volume of 200 µL in PBS. The transfection mixture was allowed to incubate at room temperature for 15 min then added dropwise at 200 µL per well to HEK293 cells. The culture was incubated at 37 °C, 5% CO_2_, under gentle agitation for 48 h. At 48 h post-transfection, 500 µL of HEK293 cells were collected into a 1.6-mL Eppendorf tube and centrifuged at 600 *× g* for 10 min at room temperature. The supernatant was discarded, and the cell pellet was lysed with 100 µL M-PER cell lysis reagent (cat # 78503, ThermoFisher) and incubated at room temperature for 10 min under gentle mixing, then 1 µL of Benzonase (cat # 70746-3, Merck) was added to the lysate and incubated on ice for an additional 10 min. Total protein concentration of lysate was determined using a Bradford assay kit (cat # 500-0006, Bio-Rad). Lysates were stored at -80 °C until used.

#### Western blot protocol

Cell lysate was thawed and diluted with 2xLDS (cat # B0008) at 1:1 ratio then heated at 75 °C for 5 min. Samples were loaded into a 12-well 4-12% Bis-Tris gel (cat # NW04122BOX, Invitrogen) and separated by SDS-PAGE in MES running buffer (cat # B0002, Invitrogen) at 200 V for 25 min. Proteins were then transferred to 0.2 µm nitrocellulose membrane (cat # 1620112, Bio-Rad) via Trans-Blot® SD Semi-Dry Electrophoretic Transfer Cell (cat # 1703940, Bio-Rad) at 200 mA for 1 h. All blots were blocked in 5% milk in TBS-T (cat # AAJ77500K8, Thermo Scientific Chemicals), then incubated overnight at 4 °C with primary antibodies (MW8, CHDI-90000942, 0.4 µg/mL, 1:10,000; 1B12, 0.4 µg/mL, 1:5,000; 11G2, 0.4 µg/mL, 1:5,000). The membranes were washed with TBS-T three times, 5 min each. After washing, the membranes were incubated at 25 °C for 30 min with either Goat anti-rabbit 800CW (cat # 92632211, Li-Cor) or Goat anti-mouse 800CW (cat # 92632210, Li-Cor) with dilution at 1:15,000. The membranes were washed with TBS-T three times, 5 min each, before scanning via Li-Cor Odyssey CLX with 700 and 800 nm channel.

#### Filter trap protocol

Cell lysate (50 µg) was diluted with PBS to a final volume of 70 µL (∼0.36 µg/µL). Samples were loaded into pre-wet cellulose acetate membranes (cat # CA023001, Sterlitech). The membranes were vacuum filtered on a Bio-DOT^TM^ apparatus (cat # 1706545, Bio-Rad). Membranes were washed with PBS twice, 5 min each, then blocked with Li-Cor Intercept blocking buffer (cat # 927-60001, Li-Cor) for 1 h at 25 °C. Membranes were incubated with primary antibodies 4C9 (0.32 µg/mL; 1:10,000), MW8 (0.37 µg/mL; 1:10,000), 1B12 (0.60 µg/mL; 1:5,000) or 11G2 (0.55 µg/mL; 1:5,000) at 4 °C overnight with gentle rocking. Membranes were washed with TBS-T three times, 5 min each, then incubated at 25 °C for 1 h with either Goat anti-rabbit 800CW (cat # 92632211, Li-Cor) or Goat anti-mouse 800CW (cat # 92632210, Li-Cor) with dilution at 1:15,000. Membranes were washed with TBS-T three times, 5 min each, and imaged via Li-Cor Odyssey CLX using the 800 nm channel.

### Characterization by immunocytochemistry

#### Transfection protocol

HEK293T cells were grown in DMEM medium supplemented by 10% FBS and 1% P/S and kept at 37 °C with 5% CO_2_ in a humidified incubator. Cells were then seeded in 60 mm dishes and transfected the following day using jetPRIME (cat # 101000015, Sartorius) with 2 µg DNA and 4 µL transfection reagent (µg DNA/ µL Reagent 1:2). After approximately 16 h, transfected cells were detached, counted and re-seeded in 96-well plates (cat # 655090, Greiner) pre-coated with fibronectin (cat # F1141 Sigma-Aldrich, 10 µg/mL in PBS, 2 h at 25 °C) at a density of 50,000 cells per well.

#### Immunofluorescence imaging

After 48 h transfection, cells were fixed in 4% paraformaldehyde for 10 min at room temperature. After three washes in PBS, cells were stored at 4 °C or processed for immunofluorescence imaging. Cells were permeabilized by 0.1% Triton X-100, blocked by 3% bovine serum albumin for 1 h at room temperature and incubated overnight with MW8 (CHDI-90000942), 1B12, and 11G2 at 1:500 dilution. After three washes in PBS, anti-rabbit Alexa Fluor ^TM^ Plus 488 (cat # A32790, ThermoFisher) for the detection of 1B12 and 11G2 and anti-mouse Alexa Fluor ^TM^ Plus 647 (cat # A32787TR, ThermoFisher) for the detection of MW8 were used as secondary antibodies at 1:1000 dilution for 1 h at room temperature. Nuclei were stained by adding Hoechst 33342 reagent (1 µg/mL, Thermo 62249) for 5 min at room temperature. Antibody-stained cell images were acquired on the Opera Phenix Plus high-content microscope (Revvity) using a 40x objective. Image analysis was performed using Signals Image Artist v. 1.3.77 software (Revvity) applying pre-defined building blocks for nuclei, cytoplasm, and aggregates segmentation.

### WB and MS analysis of proteolytic fragments

#### Proteolysis

For Arg-C digestion, 0.4 mg/mL recombinant FL Q23 HTT or FL Q48 HTT was incubated with 4 µg/mL Arg-C (cat # V1881, Promega) in a reaction buffer containing 50 mM Tris pH 7.6, 200 mM NaCl, 5 mM CaCl_2_, 2 mM EDTA, 5 mM DTT at 37 °C (0-24 h for western blot analysis, 2 h for immunoprecipitation). The reaction was stopped by adding Leupeptin (cat #78435, Thermo Scientific) to a final concentration of 184 µg/mL. For Lys-N digestion, 0.4 mg/mL recombinant FL Q23 HTT or FL Q48 HTT was incubated with 2 µg/mL Lys-N (cat # L101, IPA Therapeutics) in TBS at 37 °C for 0-24 h. The digestion was stopped by adding 500 mM 1,10-phenantroline to a final concentration of 5 mM. The digested samples were flash frozen at -80 °C before immunoprecipitation or western blot analysis.

#### Western blot protocol

Protein samples (digested and controls) were mixed with 10x SDS loading dye, [250 mM Tris (cat # T60040-5000.0, RPI), pH 6.8, 20% SDS (cat # J18220.A7, Thermo Scientific Chemicals), 10% β-mecaptomethanol (cat # M3148, Sigma Life Science), 60% glycerol (cat # BP229, Fisher Bioreagent), 0.4% bromophenol blue (cat # 11439, Sigma-Aldrich)], and heated at 70 °C for 10 min before loading on 4-12% Bis-Tris gels (cat # NW04122BOX, Invitrogen), and separated by SDS-PAGE in MES running buffer (cat # B0002, Invitrogen) at 200 V for 25 min. The proteins were transferred to nitrocellulose membrane (cat # 1620112, Bio-Rad) via Trans-Blot® SD Semi-Dry Electrophoretic Transfer Cell (cat # 1703940, Bio-Rad) at 200 mA for 1 h. All blots were blocked in 5% milk in TBS-T (cat # AAJ77500K8, Thermo Scientific Chemicals) at room temperature for 2 hrs, and then incubated with primary antibodies at 1:3,000 dilution at 4 °C overnight. The membranes were washed with TBS-T three times, 5 min each. After washing, the membranes were incubated at 25 °C for 30 min with either Goat anti-rabbit IgG (cat # 926-32211, Li-Cor) for 11G2 and 1B12, or Goat anti-mouse IgG (cat # 926-32210, Li-Cor) for other antibodies at 1:15,000 dilution.

#### Immunoprecipitation

Activated agarose beads (ThermoFisher) were crosslinked to 3B5H10 (cat # P1874, Sigma-Aldrich) according to the user manual from AminoLink™ Plus Micro Immobilization Kit (cat # 20475, ThermoFisher). Briefly, 200 µg antibody 3B5H10 was buffer exchanged to PBS using 10 kDa concentrators (cat #UFC501024, Millipore Sigma). 100 µL AminoLink^TM^ coupling resin was mixed with 3B5H10 and 6 µL 5 M cyanoborodydride in 400 µL coupling buffer to crosslink at 25 °C for 90 min. The crosslinked beads were quenched with quenching buffer and washed extensively with PBS and stored at 4 °C. To perform immunoprecipitation, 30 µL 3B5H10 crosslinked beads and control beads (provided in the kits) were first washed with 500 µL wash buffer containing PBS and 0.1% w/v CHAPS. 500 µL Arg-C or Lys-N digested FL Q48 HTT was incubated with 3B5H10 crosslinked beads and control beads for 1 h at 4 °C. The beads were washed 1 x 500 µL wash buffer followed by 2 x 500 µL PBS. The bound proteins were eluted with 2 x 50 µL 0.5% RapiGest (cat # 186001860, Waters) followed by 50 µL 0.2% w/v SDS at 70 °C for 2 min. The eluted peptides were flash frozen at -80 °C before MS or western blot analysis.

#### Mass Spectrometry analysis

Samples eluted from 3B5H10 cross-linked beads (40 µL) were acidified by adding TFA (0.5% v/v) at 37 °C for 30 min to degrade RapiGest. The samples were then centrifuged at 17,000 × *g* for 10 min at 4 °C to remove the insoluble materials. The supernatants were diluted with 40 µL water and filtered with 0.2 µM spin filter (cat # UFC30LG25, Millipore Sigma) before loading onto the LC-MS (Shimadzu LC-20AR; SCIEX TripleTOF 6600) at 20 µL per injection. The peaks were separated using a ProSwift RP-4H HPLC column (cat # 0666640, ThermoFisher) with a 10 min 5%-65% gradient of mobile phase B (mobile phase A - H_2_O, 0.09% formic acid and 0.02% TFA; mobile phase B – 80% isopropanol, 20% acetonitrile, 0.09% formic acid and 0.02% TFA). The intact mass was analyzed using BioPharmaView 2.0 Intact Mass module from SCIEX.

## Results

### Immunization and clonal selection strategy

The immunization and screening strategy was designed around a region of homology between the human and mouse HTT sequence spanning A83–K99 (Fig 1, dashed box). By raising antibodies against peptides within this region, we reasoned that we could identify not only clones specific to the engineered C- or N-terminal neoepitopes, but also clones that happened to cross-react with the native, full-length mouse and human protein—expanding the utility of the resulting antibody panel for preclinical use.

The immunizing peptides accordingly contained the free C-terminal HTTexon1 P90 carboxylic acid (AEEPLHRP-OH) or the free N-terminal HTTexon2 K91 amine (NH2-KKELSATK). These same peptides served as positive-screening antigens by ELISA, with counter-screening peptides spanning shorter and longer flanking fragments used to define epitope specificity (Fig 1, solid box). Following FACS and ELISA screening of B-cell culture supernatants (ABclonal, Woburn, MA), a subset of clones was selected for recombinant expression, purification, and full validation.

From the HTTexon1 immunization, C-terminal P90 neoepitope-specific clones 1B12 and 11G2 were advanced to further validation. As anticipated, additional clones emerged that reacted across all Exon1 counter-screening peptides; among these, clones 27F5 and 31C10 showed robust reactivity with both human and mouse FL HTT. These PRD-directed, species cross-reactive mAbs represent the added benefit of this strategy, expanding the available antibody repertoire for preclinical HTT detection assays.

The HTTexon2 immunization, in contrast, yielded N-terminal K91 neoepitope-specific clones but none with ELISA reactivity to the Exon2 counter-screening peptides—indicating that this strategy did not produce mouse-human FL HTT cross-reactive clones. Of the HTTexon2 N-terminal neoepitope-specific mAbs identified, clone 7G10 was selected for further validation.

### Characterization of mAbs by western blotting and surface plasmon resonance analysis with recombinant proteins

The selected recombinant mAb clones were characterized by WB and SPR using recombinant protein standards containing the relevant neoepitopes, HTT Q23 Exon1 and HTT 91-171 C-term MBP, in comparison to FL HTT. To ensure that the HTTexon1 P90 and HTTexon2 K91 protein standards contained the requisite neoepitopes, they were produced using a SUMO (Small Ubiquitin-like Modifier) fusion tag cleaved by the SUMO-specific protease ULP1 (Ubl-specific protease 1), which reveals the neoepitope cleanly, without residual amino acids or miscleavage products [17]. The selectivity profiles of the newly generated mAbs were compared with three known antibodies recognizing similar regions of HTT: (i) mAb 4C9, a human-specific antibody with an epitope mapped to aa 51–71 within the human PRD of HTTexon1 [18]; (ii) mAb MW8 (CHDI-90000942), mapped to aa 83–91 [12], which recognizes HTTexon1 aggregates; and (iii) MAB5490 (clone 1H6), mapped to aa 115–129 within HTTexon2 [7] (Fig 2).

**Fig 2.**
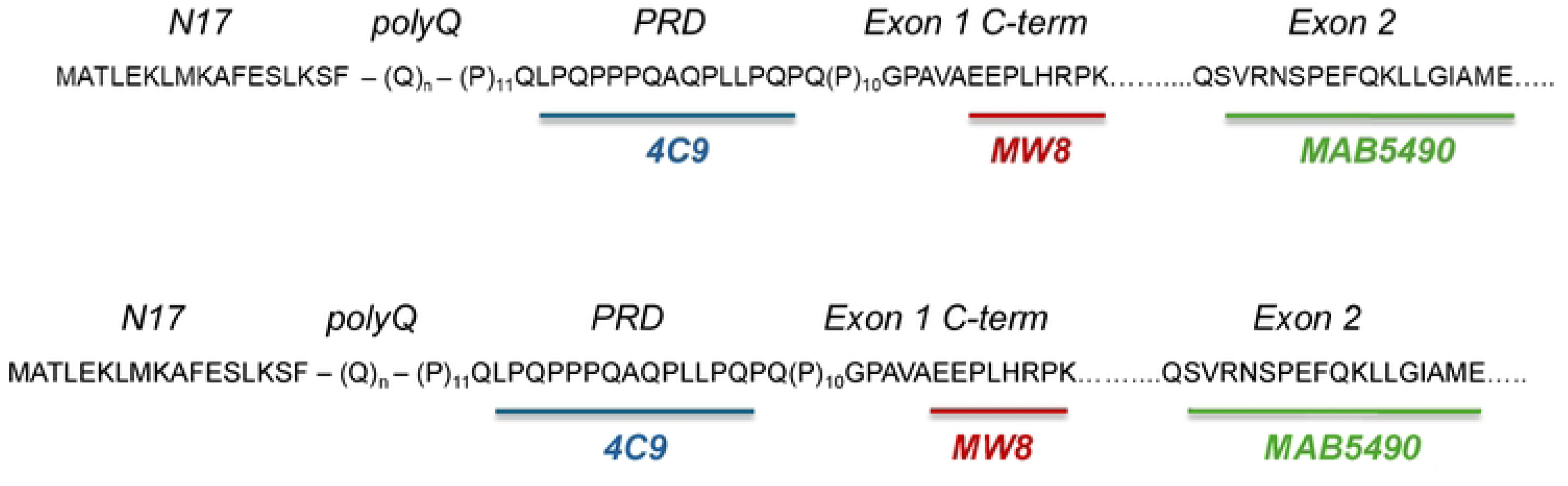
Epitope map for human HTT N-terminal comparator mAbs used in WB and SPR analysis.

In western blot analyses, mAbs MW8, 1B12, and 11G2 showed clear selectivity for recombinant human HTT Q23 Exon1 protein over human FL Q23 HTT. mAb 4C9 recognized both HTT Q23 Exon1 and FL Q23 HTT, consistent with its epitope lying within both, but did not react with the HTT 91–171 fragment, confirming that its epitope does not extend into this region. Conversely, MAB5490 detected both the HTT 91–171 fragment and FL Q23 HTT, as expected, whereas mAb 7G10 selectively recognized only the HTT 91–171 fragment bearing the N-terminal K91 neoepitope (Fig 3). The appearance of two bands for the HTT 91-171 standard at MW 50 kDa and 100 kDa in Fig 3 is due to the propensity of this recombinant protein standard to form a stable dimer.

**Fig 3.**
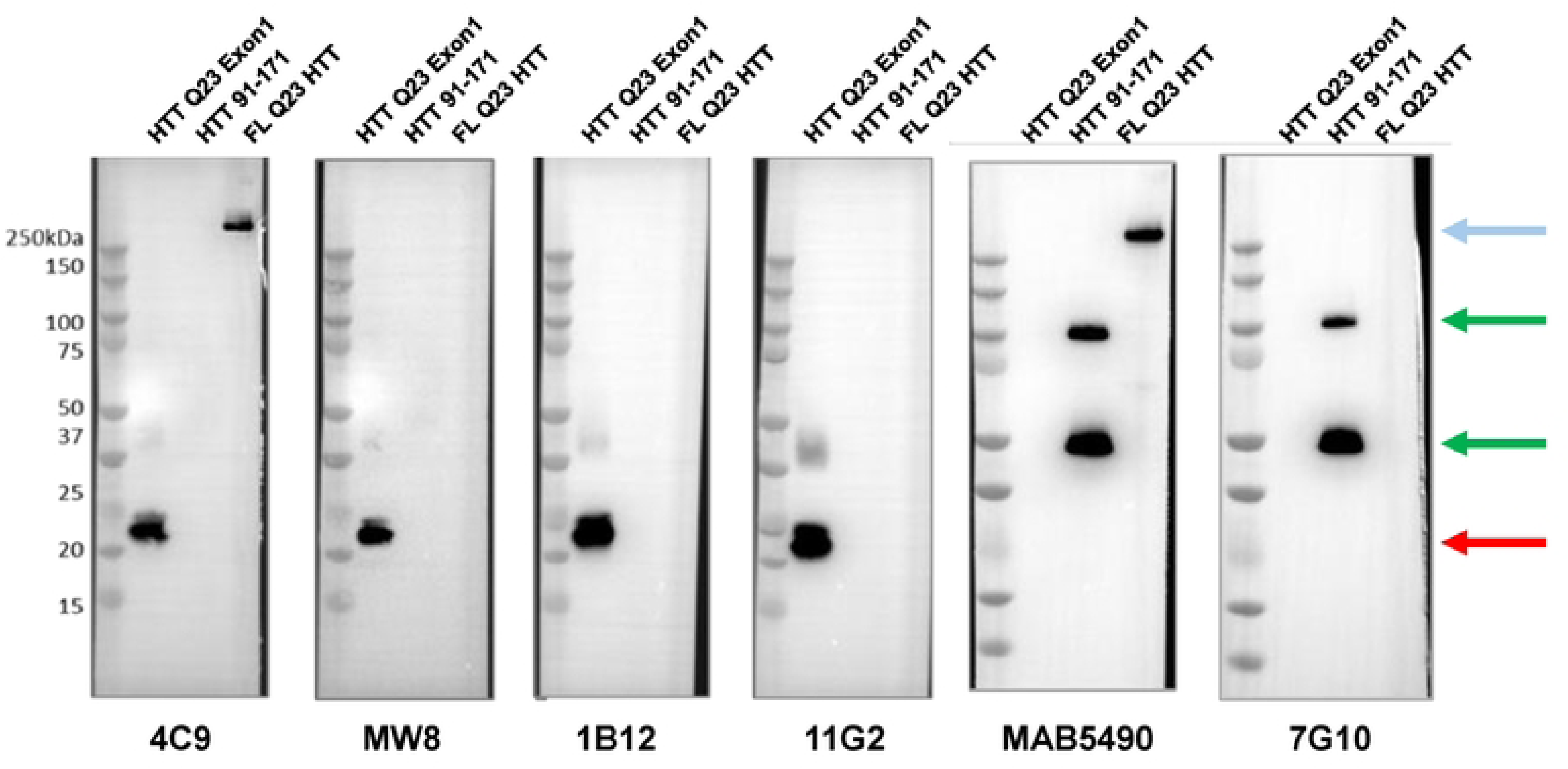
Western blot comparison for mAbs 4C9, MW8, 1B12, 11G2, MAB5490, and 7G10. Lanes: (1) protein standard ladder; (2) HTT Q23 Exon1, human; (3) HTT (91-171) C-term MBP; (4) FL Q23 HTT, human. Blue arrow indicates FL HTT. Green arrows indicate HTT (91-171) C-term MBP monomer/dimer. Red arrow indicates HTT Q23 Exon1.

Clones 27F5 and 31C10 also showed strong recognition of mouse FL Q7 HTT, in contrast to the human-PRD-selective mAb 4C9, which did not react with mouse FL Q7 HTT. mAb MAB2166, with an epitope mapped to aa 443–457 [19] outside the PRD and conserved between species, served as a comparator to confirm that mouse FL Q7 HTT was present and detectable on the blot independent of PRD sequence divergence (Fig 4). The human-mouse cross-reactive PRD mAbs 27F5 and 31C10 detected both human and mouse FL HTT as well as the human HTT Q23 Exon1 fragment (Fig 4).

**Fig 4.**
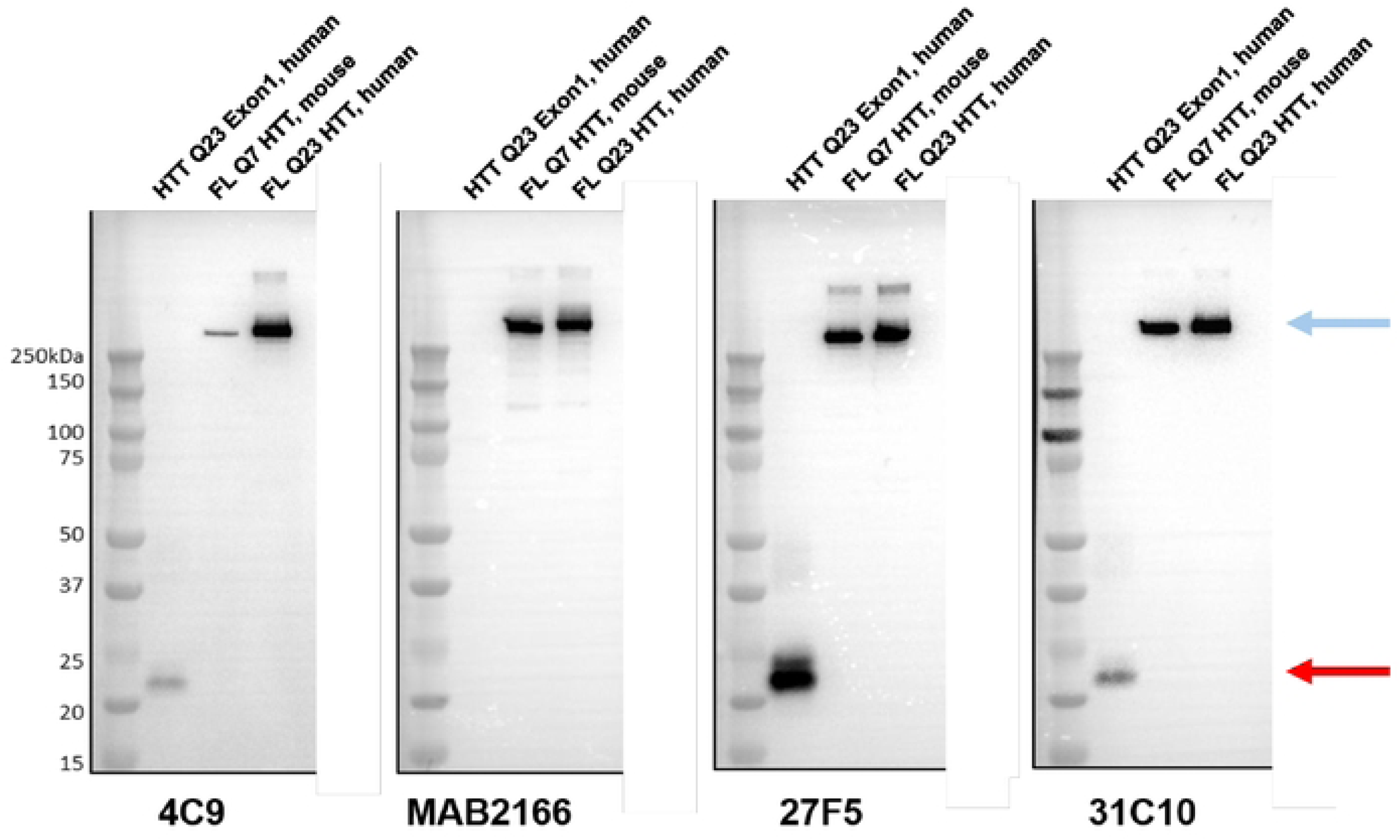
Western blot comparison of mAbs 4C9, MAB2166, clone 27F5 and 31C10. Lanes: (1) protein standard ladder; (2) HTT Q23 Exon1, human; (3) FL Q7 HTT, mouse; (4) FL Q23 HTT, human. Blue arrow indicates FL HTT (mouse and human). Red arrow indicates HTT Q23 Exon1.

The relative binding affinity kinetics were determined for the mAbs by SPR and results are summarized in Table 1. SPR revealed similar affinities for mAbs 4C9, 1B12, and 11G2 to the human HTTexon1 protein standard (KD values of 76 ± 28 nM, 25 ± 10 nM, and 30 ± 12 nM, respectively), with significantly higher affinity than MW8 (KD = 761 ± 49 nM). Consistent with the western blot data, MW8, 1B12, and 11G2 showed no detectable binding to FL Q23 HTT, whereas mAb 4C9 bound FL Q23 HTT with high affinity (KD = 17 ± 4 nM) (Table 1). Representative sensorgrams for 4C9, MW8, 1B12, and 11G2 against HTT Q23 Exon1 compared to FL Q23 HTT are shown in Fig 5. The human-mouse cross-species reactive mAbs 27F5 and 31C10 showed high affinity for mouse FL Q7 HTT (27F5 KD = 10 ± 2 nM; 31C10 KD = 13 ± 2 nM) and human FL Q23 HTT (27F5 KD = 8 ± 2 nM; 31C10 KD = 10 ± 0.1 nM). The N-term K91 neoepitope mAb 7G10 bound its cognate protein standard with high affinity (KD = 2 ± 0.6 nM) and showed no detectable binding to human FL Q23 HTT (Table 1).

**Fig 5.**
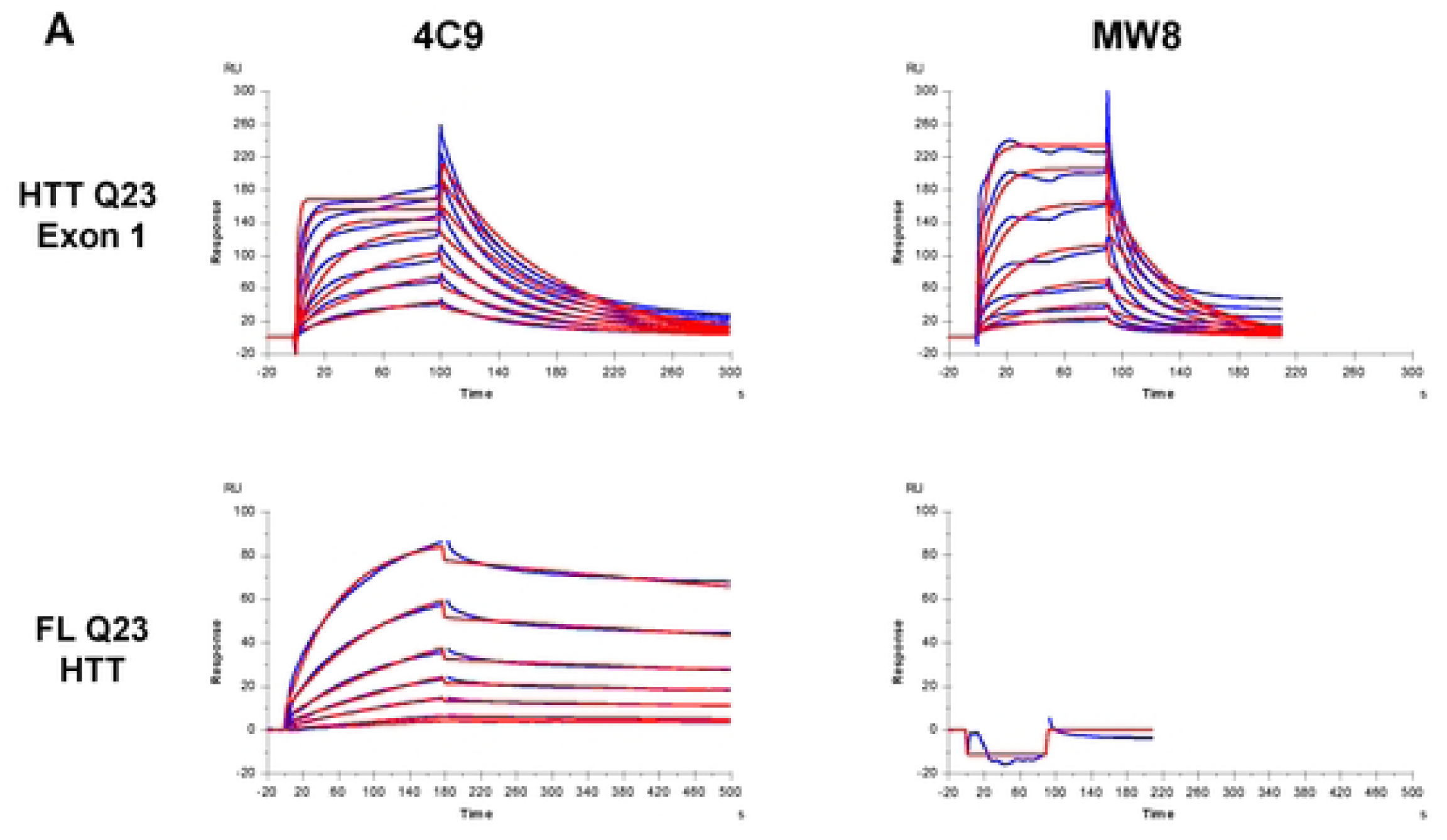
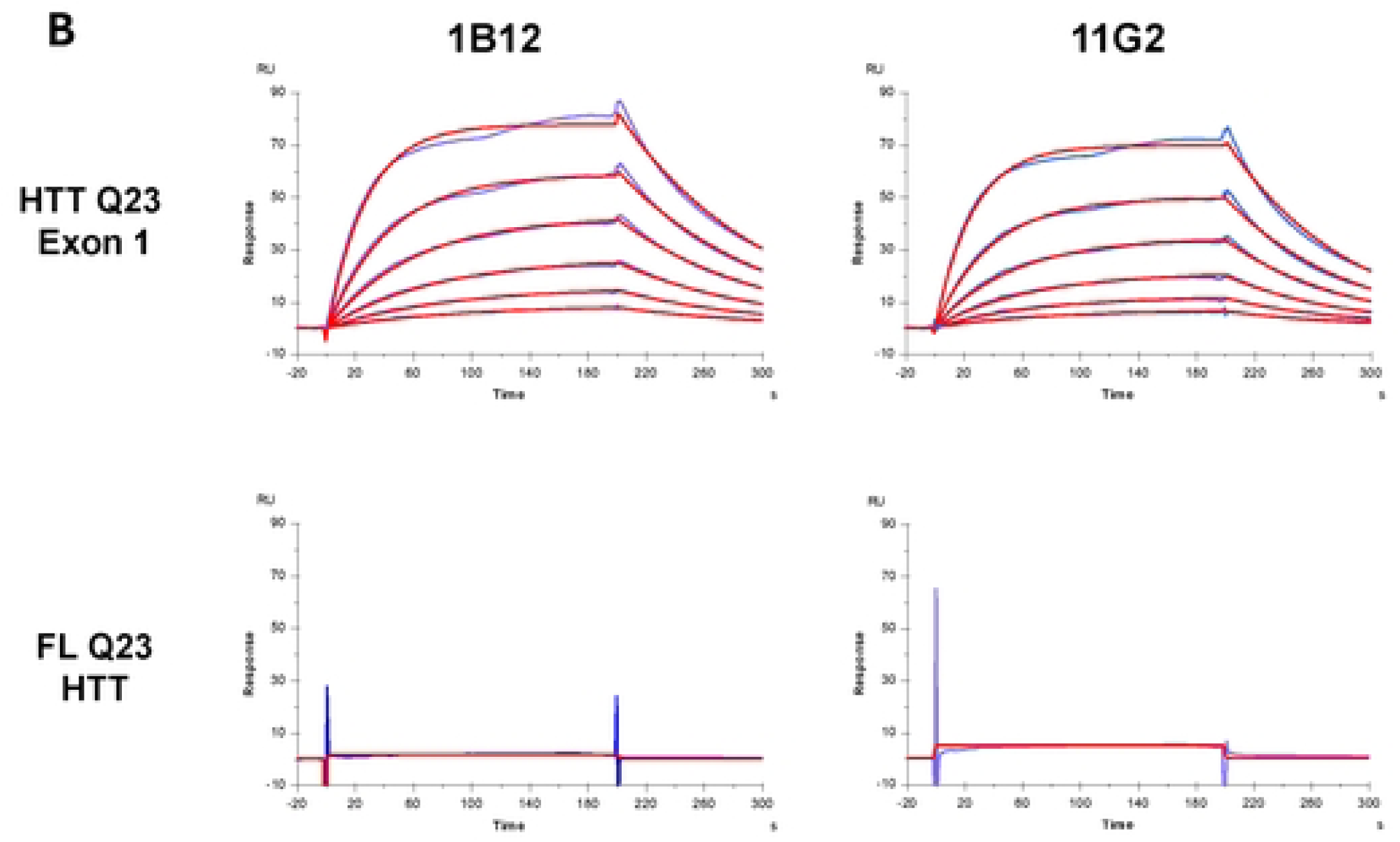
Sensorgrams (blue) and fitting (red) of antibodies interactions with HTT proteins: (A) mAb 4C9 and MW8 interaction with HTT Q23 Exon1 compared to FL Q23 HTT, human; (B) P90 mAbs 1B12 and 11G2 interaction with HTT Q23 Exon1 compared to FL Q23 HTT.

**Table 1:** SPR binding kinetics for mAbs 4C9, MW8, 1B12, 11G2 against proteins FL Q23 HTT and HTT Q23 Exon1; 27F5 and 31C10 against FL Q7 HTT (mouse) and FL Q23 HTT (human); 7G10 against HTT 91-171 C-term MBP and FL Q23 HTT. Association (*k_on_*) and dissociation (*k_off_*) rates and affinity (K_D app_) reported as mean ± standard deviation of individual experiments (n).

| mAb clone | Protein | $k_{on}$ ($M^{-1}s^{-1}$) | $k_{off}$ ($s^{-1}$) | $K_{D\ app}$ (nM) | n |
| --- | --- | --- | --- | --- | --- |
| 4C9 | HTT Q23 Exon1 | $2 \pm 1 \times 10^5$ | $1.5 \pm 0.2 \times 10^{-2}$ | $76 \pm 28$ | 2 |
| 4C9 | FL Q23 HTT, human | $4.5 \pm 0.5 \times 10^4$ | $7 \pm 2 \times 10^{-4}$ | $17 \pm 4$ | 3 |
| MW8 | HTT Q23 Exon1 | $5.5 \pm 0.6 \times 10^4$ | $1.8 \pm 0.3 \times 10^{-2}$ | $761 \pm 49$ | 4 |
| MW8 | FL Q23 HTT, human | No binding |  |  | 2 |
| 1B12 | HTT Q23 Exon1 | $4 \pm 1 \times 10^5$ | $1 \pm 0.6 \times 10^{-3}$ | $25 \pm 10$ | 3 |
| 1B12 | FL Q23 HTT, human | No binding |  |  | 3 |
| 11G2 | HTT Q23 Exon1 | $3.5 \pm 0.8 \times 10^5$ | $1 \pm 0.2 \times 10^{-2}$ | $30 \pm 12$ | 3 |
| 11G2 | FL Q23 HTT, human | No binding |  |  | 3 |
| 27F5 | FL Q7 HTT, mouse | $3.2 \pm 0.4 \times 10^4$ | $3.2 \pm 0.2 \times 10^{-4}$ | $10 \pm 2$ | 2 |
| 27F5 | FL Q23 HTT, human | $4 \pm 1 \times 10^4$ | $3.3 \pm 0.1 \times 10^{-4}$ | $8 \pm 2$ | 2 |
| 31C10 | FL Q7 HTT, mouse | $2.66 \pm 0.08 \times 10^4$ | $3.4 \pm 0.3 \times 10^{-4}$ | $13 \pm 2$ | 2 |
| 31C10 | FL Q23 HTT, human | $3.5 \pm 0.5 \times 10^4$ | $3.3 \pm 0.4 \times 10^{-4}$ | $10 \pm 0.1$ | 2 |
| 7G10 | HTT 91-171, C-term MBP | $2.6 \pm 0.4 \times 10^5$ | $5 \pm 1 \times 10^{-4}$ | $2 \pm 0.6$ | 4 |
| 7G10 | FL Q23 HTT, human | No binding |  |  | 4 |

We next assessed binding of the Exon1-directed mAbs to pre-formed fibrils of HTT Q43 Exon1, prepared according to Reif et al. [17]. Prior to SPR analysis, fibrils were sonicated to generate a more homogeneous suspension, then immobilized on CM5 sensor chips via amine coupling. The sensorgrams are shown in Fig 6. The relative binding kinetics were determined using a bivalent fitting model and are summarized in Table 2. Under these conditions, mAbs 4C9, 1B12, and 11G2 displayed comparable affinities for HTT Q43 Exon1 fibrils (KD = 0.5 ± 0.1 nM, 0.2 ± 0.03 nM, and 0.5 ± 0.2 nM, respectively). In contrast, MW8 exhibited a significantly faster dissociation rate, resulting in a weaker apparent affinity (KD = 12 ± 3 nM) (Table 2).

**Fig 6.**
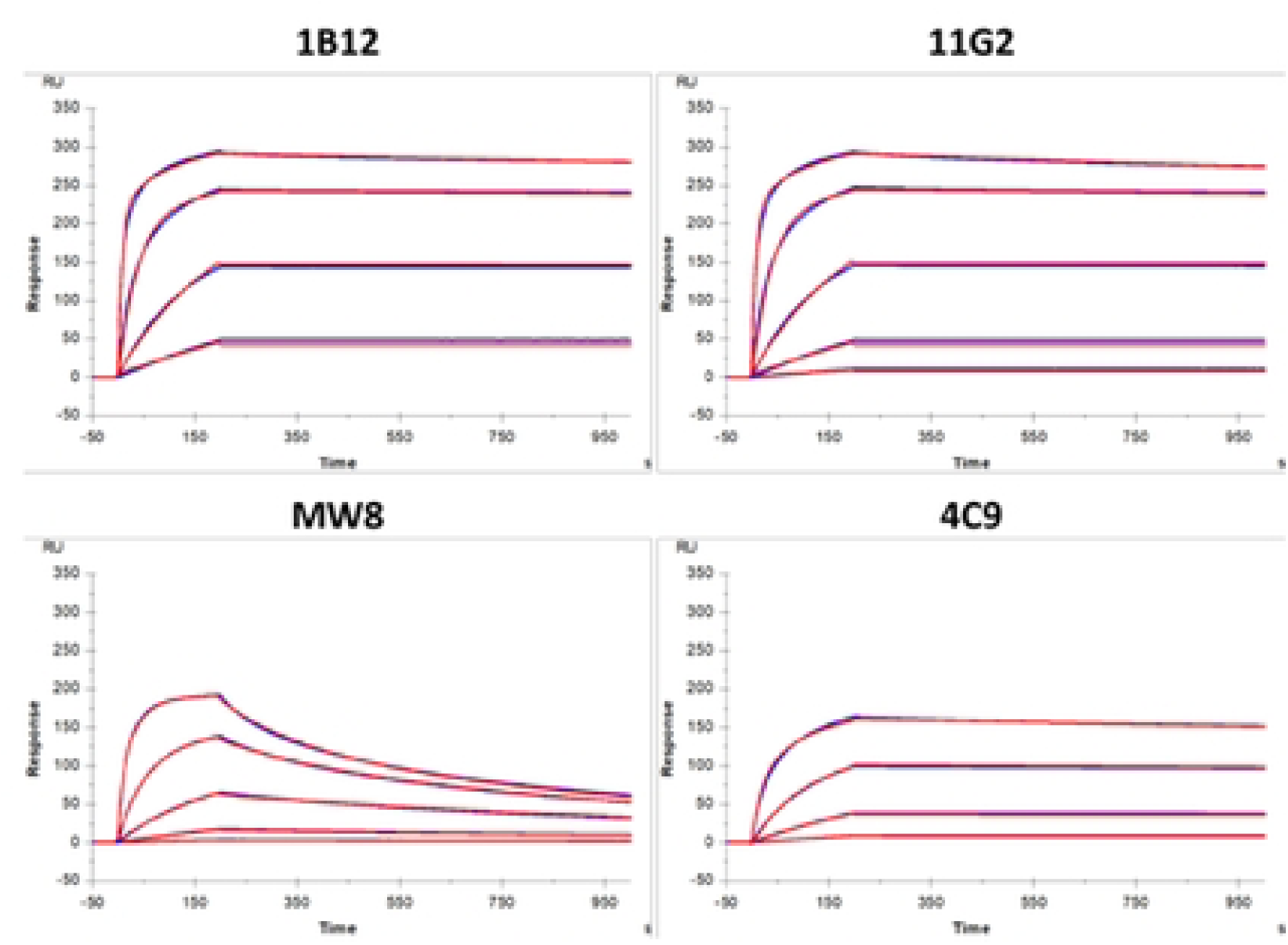
Sensorgrams (blue) and fitting (red) for 1B12, 11G2, MW8 and 4C9 interactions with HTT Q43 Exon1 fibrils.

**Table 2:**
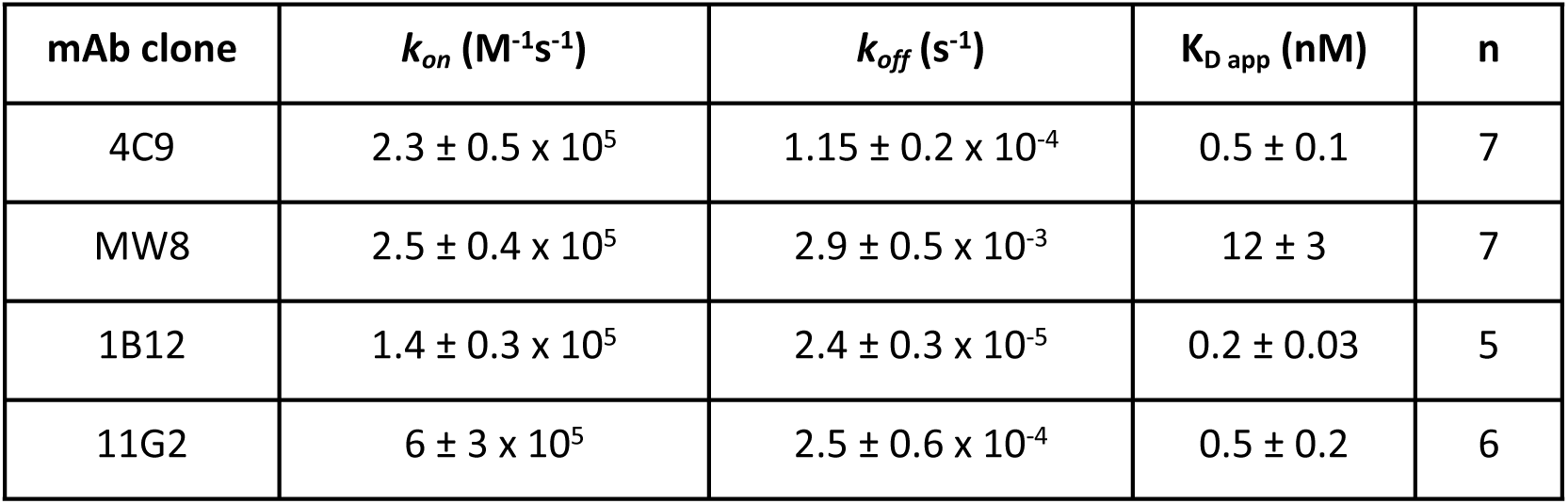
SPR binding kinetics for for mAbs 4C9, MW8, 1B12, 11G2 interaction with HTT Q43 Exon1 fibrils. Association (*k_on_*) and dissociation (*k_off_*) rates and affinity (K_D app_) reported as mean ± standard deviation of individual experiments (n).

| mAb clone | $k_{on}$ ($M^{-1}s^{-1}$) | $k_{off}$ ($s^{-1}$) | $K_{D\ app}$ (nM) | n |
| --- | --- | --- | --- | --- |
| 4C9 | $2.3 \pm 0.5 \times 10^5$ | $1.15 \pm 0.2 \times 10^{-4}$ | $0.5 \pm 0.1$ | 7 |
| MW8 | $2.5 \pm 0.4 \times 10^5$ | $2.9 \pm 0.5 \times 10^{-3}$ | $12 \pm 3$ | 7 |
| 1B12 | $1.4 \pm 0.3 \times 10^5$ | $2.4 \pm 0.3 \times 10^{-5}$ | $0.2 \pm 0.03$ | 5 |
| 11G2 | $6 \pm 3 \times 10^5$ | $2.5 \pm 0.6 \times 10^{-4}$ | $0.5 \pm 0.2$ | 6 |

Taken together, these binding data show that the P90 neoepitope-specific mAbs 1B12 and 11G2 recognize HTTexon1 with high affinity in both its monomeric, disaggregated form and as fibrillar aggregates; that the K91 neoepitope mAb 7G10 distinguishes the HTT 91–171 fragment from FL HTT; and that the cross-reactive mAbs 27F5 and 31C10 recognize FL HTT in both species as well as the human HTTexon1 fragment.

### Validation of P90 mAb fragment-length specificity by western and filter-trap immunoblotting

In an experiment analogous to that reported by Landles et al. [4], we next assessed the N-terminal HTT fragment-length specificity of mAbs 1B12 and 11G2 in comparison to MW8 (CHDI-90000942). Antibody reactivity was evaluated against a series of HTT fragments terminating at L87, R89, P90, K91, E93, S95, and K99 (Fig 7). Corresponding DNA constructs were transiently expressed in HEK cells, and cell lysates were analyzed by western blotting. Recombinant human HTTexon1 Q23 and Q43 proteins were included as positive controls and confirmed that the overexpressed fragments migrated at the expected molecular weights (Fig 7).

**Fig 7.**
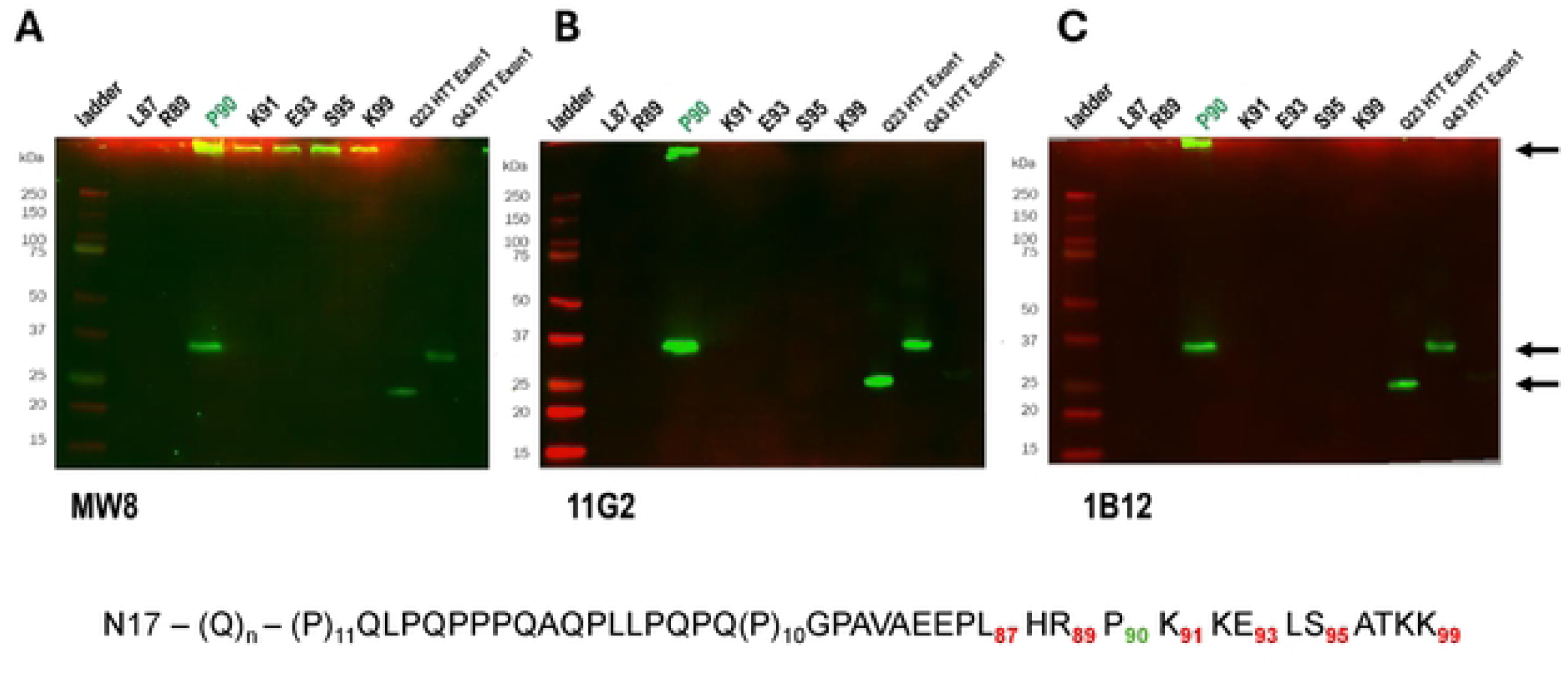
Western blot of (A) MW8 (CHDI-90000942), (B) 11G2, and (C) 1B12 antibodies against a series of N-term HTT fragments of varying length overexpressed in HEK cells. Arrows indicate aggregate signal (top), HTTexon1 1-90 fragments (lower). The protein sequence indicating fragment lengths is shown below. Lanes: (1) ladder; (2) C-term L87; (3) C-term R89; (4) C-term P90; (5) C-term K91; (6) C-term E93; (7) C-term S95; (8) C-term K99; (9) recombinant HTT Q23 Exon1; (10) recombinant HTT Q43 Exon1.

As expected, mAbs 1B12, 11G2, and MW8 detected the C-terminal P90 fragment, both as a monomeric species and as higher-molecular-weight aggregates retained at the top of the gel (lane 4, Fig 7A, 7B, and 7C). In contrast to the findings of Landles et al., who did not detect aggregated fragments, our studies show that MW8 detected aggregates formed by longer fragments (K91, E93, S95, and K99; lanes 5–8, Fig 7A). Notably, mAbs 1B12 and 11G2 were selective for the C-terminal P90 fragment and did not react with monomeric or aggregated forms of the longer fragments. All blots were re-probed with the N17-directed mAb 2B7 to confirm the expression of the HTT fragments (Supporting Information, Fig S3).

The same cell lysates were next processed by filter-trap assay, which captures aggregates for direct immunoblot detection on the capture membrane (Fig 8). Results were consistent with the western blot data: MW8 detected aggregates for HTT Q48 1–90 as well as for the fragments terminating at K91, E93, S95, and K99. By filter trap, both 11G2 and 1B12 detected the HTT Q48 1–90 aggregate, with a faint additional signal observed for the HTT Q48 1–91 fragment. The human PRD mAb 4C9, used as a comparator, produced signal across all HTT Q48 fragments, with a faint signal also detected in HTT Q23 1–90-transfected and untransfected cells, likely reflecting detection of endogenous HTT trapped on the membrane.

**Fig 8.**
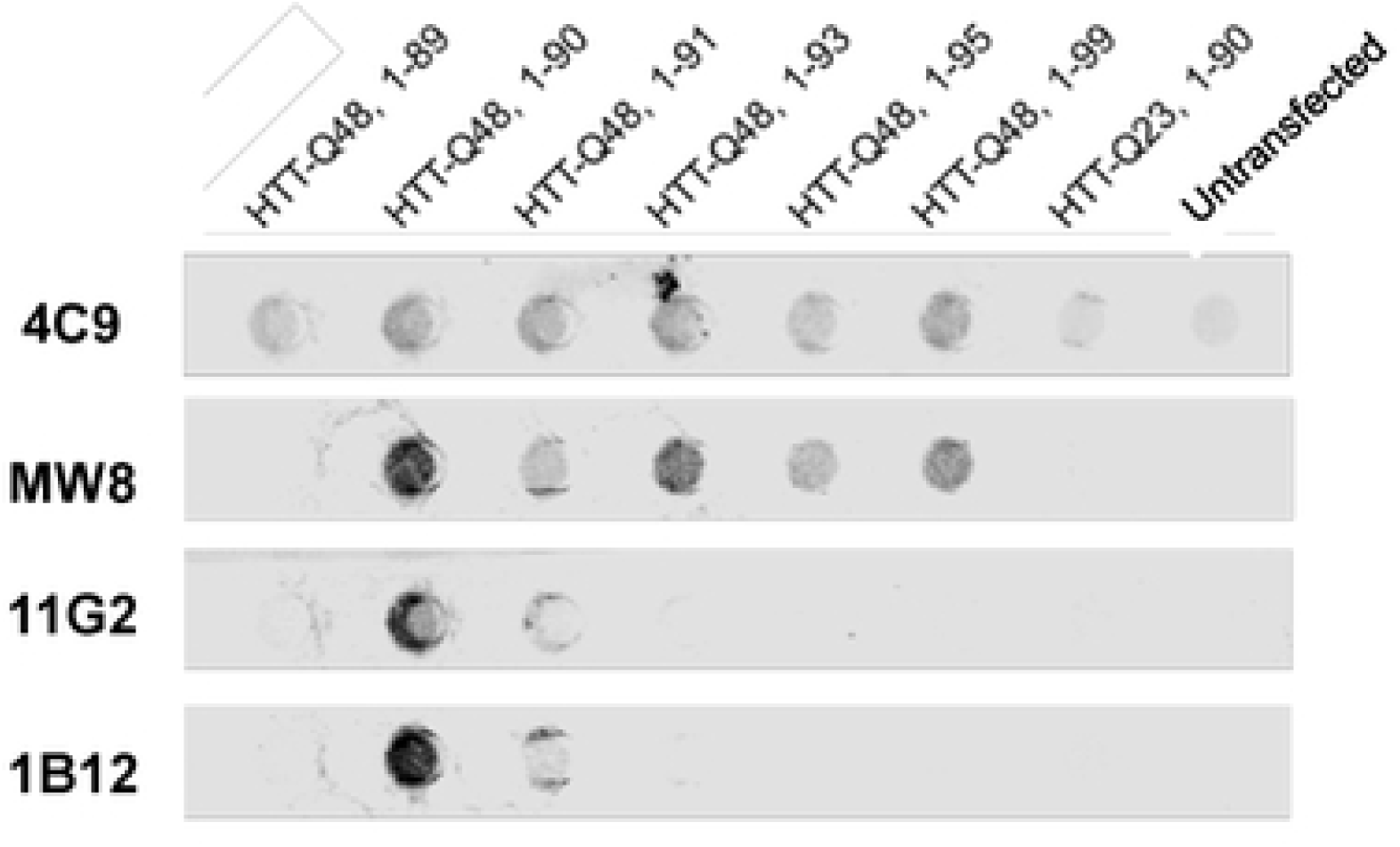
Filter-trap immunoblot of 4C9, MW8, 11G2, and 1B12 antibodies against HEK cell lysates with transiently expressed HTT N-term fragments, columns: (1) HTT Q48, 1-89; (2) HTT Q48, 1-90; (3) HTT Q48, 1-91; (4) HTT Q48, 1-95; (5) HTT Q48, 1-99; and (6) HTT Q23, 1-90. The final column is lysate from untransfected cells.

Given the striking difference in MW8 selectivity between our findings in Fig 7 and the previous report [4], we provided the Landles lab with the same plasmids to repeat the study using their own source of MW8 (Merck MABN2529) and according to their published methods. Using transiently transfected COS-1 cells, the Landles lab independently confirmed by WB (Fig 9) that the Merck-sourced MW8 shows higher C-terminal P90 neoepitope specificity than the CHDI-sourced MW8 (CHDI-90000942) used in our initial experiments (Fig 7): with the Merck-sourced antibody, only the HTT Q48 1–90 soluble and aggregated signal was detected, comparable to 11G2 and 1B12. A comparison western blot using mAb 4C9 is also shown in Fig 9, confirming the relative expression of each fragment.

**Fig 9.**
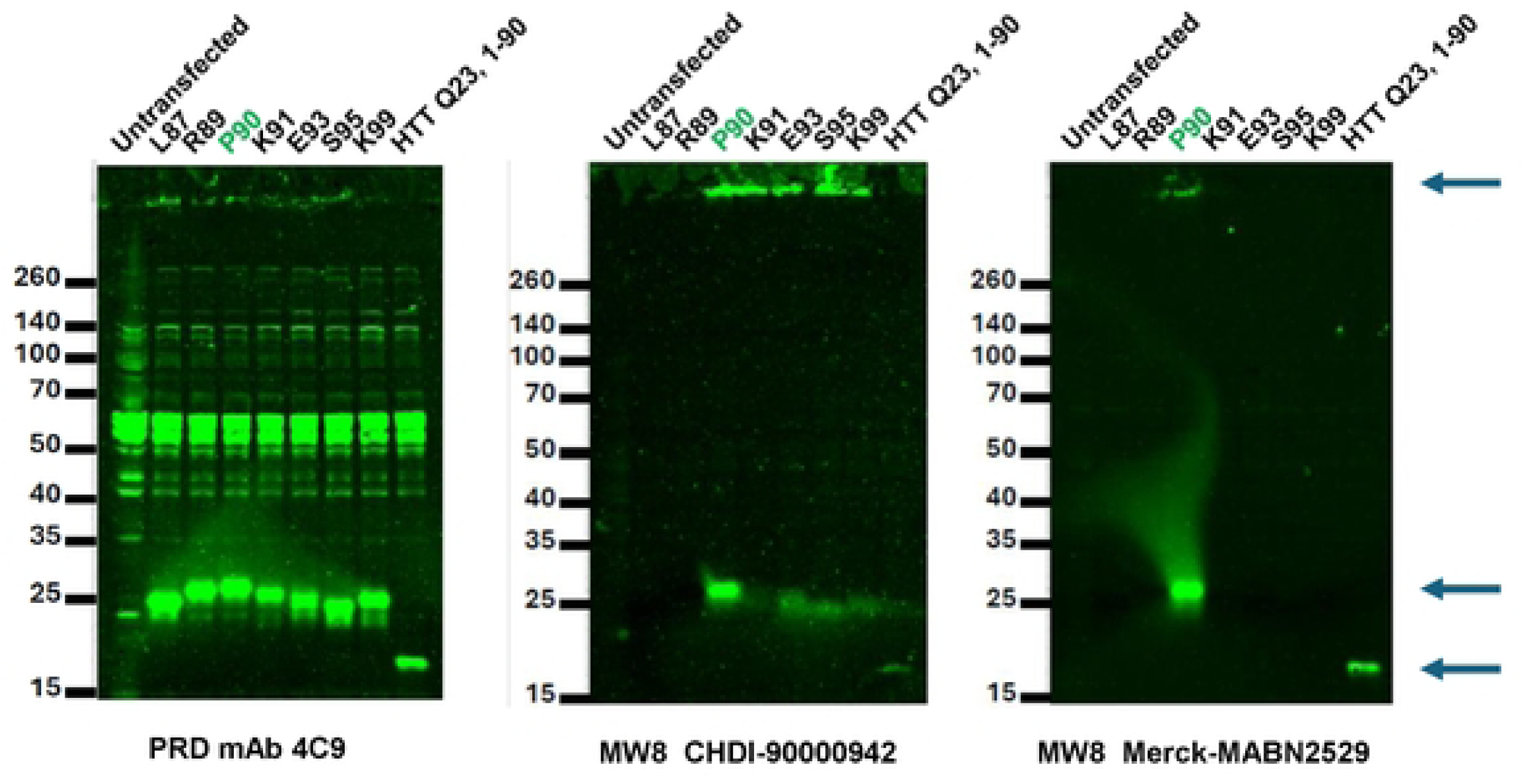
Western blot comparison of mAb 4C9, CHDI MW8 (CHDI-90000942), and Merck MW8 (MABN2529) against a series of N-term HTT fragments of varying length overexpressed in COS-1 cells. Arrows indicate aggregate signal (top); HTT Q48, 1-90 (middle); HTT Q23, 1-90 (lower). Independent replicate experiment performed by C. Landles, UCL.

### Characterization of P90 mAbs by immunocytochemistry

To correlate with the western blot results, we further characterized clones 1B12 and 11G2 by ICC using HEK293T cells overexpressing Q48 HTTexon1 (aa 1-90) protein fragments, with non-transfected cells used as a control. Given the history of MW8 use in IHC applications [20–22], we included MW8 (CHDI-90000942) as a comparator. At 1:500 dilution on fixed and permeabilized cells, both 1B12 and 11G2 showed minimal background staining in non-transfected cells (Fig 10A and B, NT, top rows) and clear signal that co-localized with MW8 in Q48 HTTexon1 (1–90)-expressing cells (Fig 10A and B, bottom rows).

**Fig 10.**
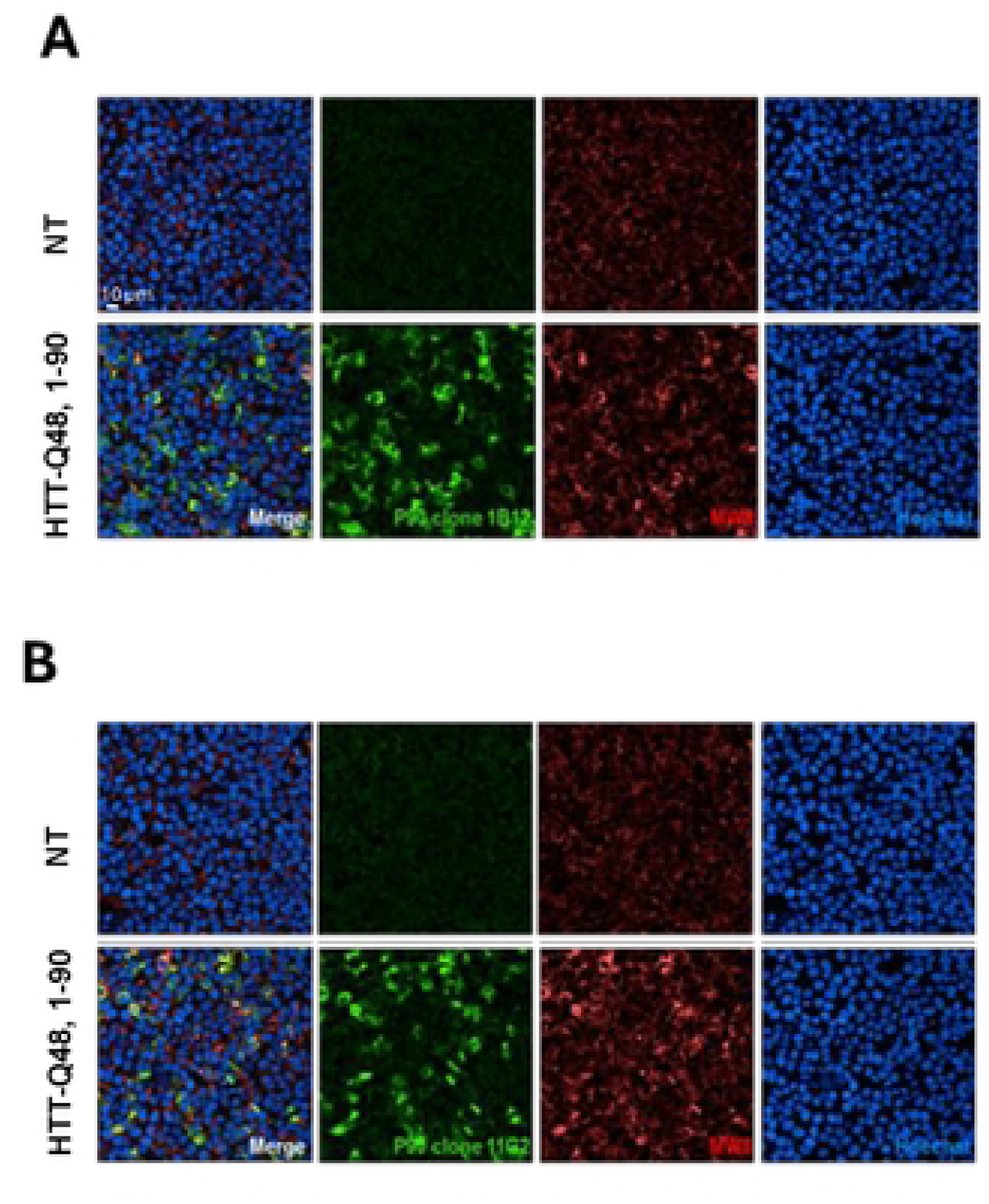
Representative images of P90 mAb 1B12 (A) and 11G2 (B) co-stained with MW8 (CHDI-90000942) in untransfected cells (NT, top row) and expressing HTT-Q48, 1-90 (bottom row). Antibodies were diluted 1:500 corresponding to 6 µg/ml for 1B12, 5.5 µg/ml for 11G2, and 7 µg/ml for MW8. Nuclear counterstain with Hoechst (far right) and merge of P90 mAb, MW8, and Hoechst stain (far left). Scale Bar: 10 µm.

We next examined the specificity of P90 mAb clone 1B12 for HTT fragments of varying length (1–89, 1–90, 1–91, and 1–93) overexpressed in HEK293 cells, again in comparison to MW8 (CHDI-90000942). Clone 1B12 detected both Q23 and Q48 HTT 1–90 overexpressed fragments, with diminished signal for HTT 1–91 (Q48) and no signal above background for the shorter (1–89) or longer (1–93) fragments (Fig 11). Consistent with the WB and filter-trap results (Figs 7 and 8), MW8 signal was observed for all fragment lengths except the truncated 1–89 fragment. Notably, aggregate staining with 1B12 co-localized with MW8-positive aggregates (Fig 11B). The P90 mAb also showed a diffuse staining pattern, consistent with detection of monomeric HTTexon1.

**Fig 11.**
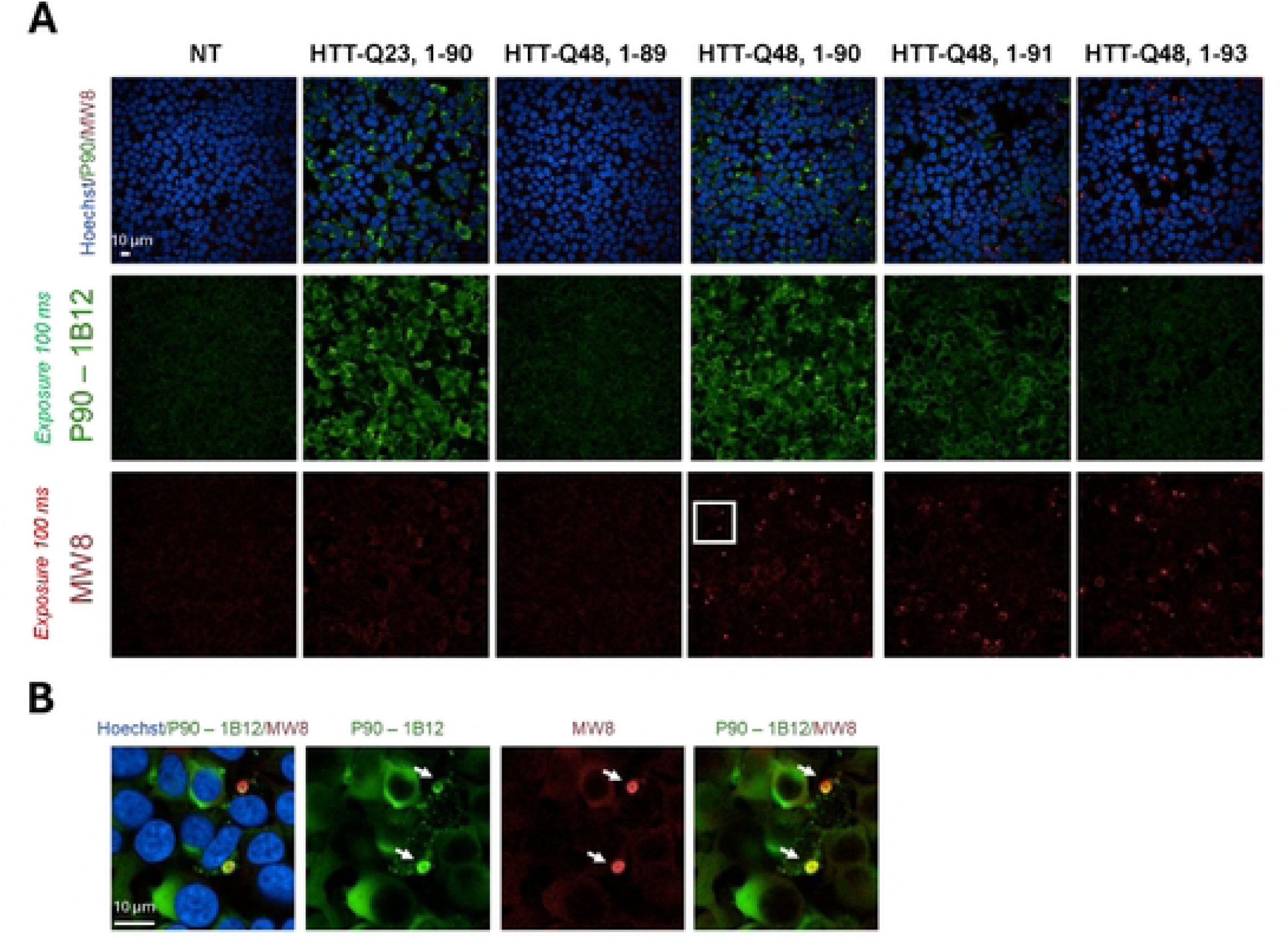
(A) Representative images of P90 clone 1B12 and MW8 (CHDI-90000942) co-staining in the indicated transfected and non-transfected cells (NT). (B) Representative images of MW8 and P90 clone 1B12 colocalization (expansion of box inset in Fig A). Aggregates are indicated by arrows. Scale bar: 10 µm.

### WB and MS validation of P90 mAb neoepitope specificity by proteolysis

To evaluate the specificity of the P90 and K91 mAbs for the putative products of proteolytic cleavage at the P90–K91 site, recombinant human FL Q23 HTT protein was digested with Lys-N or Arg-C proteases, and the resulting fragments were characterized by WB and mass spectrometry (MS). The predicted Lys-N cleavage sites [23] in FL HTT would reveal the P90 C-terminal and K91 N-terminal epitopes, among others, while Arg-C cleavage would be predicted to reveal only the R89 C-terminal and P90 N-terminal epitopes (Fig 12). In this experiment, we expected to clearly distinguish P90- and K91-mAb-reactive bands in the Lys-N digest, while the Arg-C proteolytic fragments should not be detected by either mAb.

**Fig 12.**
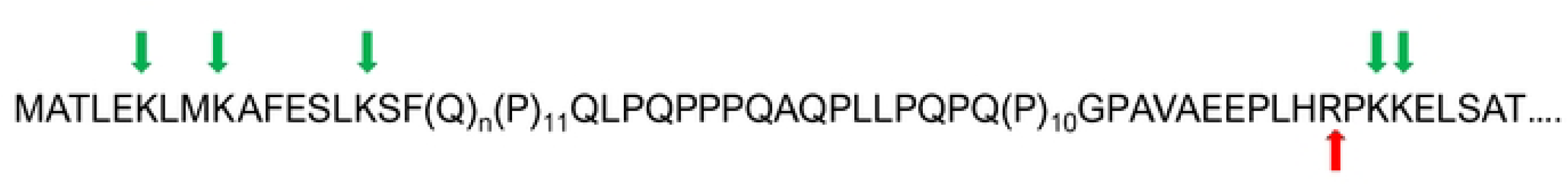
Green arrows indicate the predicted site [23] of Lys-N-term proteolytic cleavage at K6, K9, K15, K91, or K92. Red arrow indicates the predicted site of Arg-C-term proteolytic cleavage at R89.

As predicted, Lys-N digestion of FL Q23 HTT over a 24 h period (Fig 13) revealed multiple fragments detectable by mAb 4C9 (PRD epitope) and MAB5490 (Exon2 epitope, aa 115–129). The neoepitope-specific mAbs 1B12 (C-term P90) and 7G10 (N-term K91) revealed a simpler profile, detecting fewer bands over time. After complete digestion, Lys-N produced a P90-positive signal at approximately 25 kDa. This same 1B12-positive 25 kDa band was also 4C9-positive, indicating that it contains the PRD domain. The Lys-N digest also produced multiple MAB5490-positive bands, a subset of which contained the N-terminal K91 neoepitope, observed as 7G10-positive bands.

**Fig 13.**
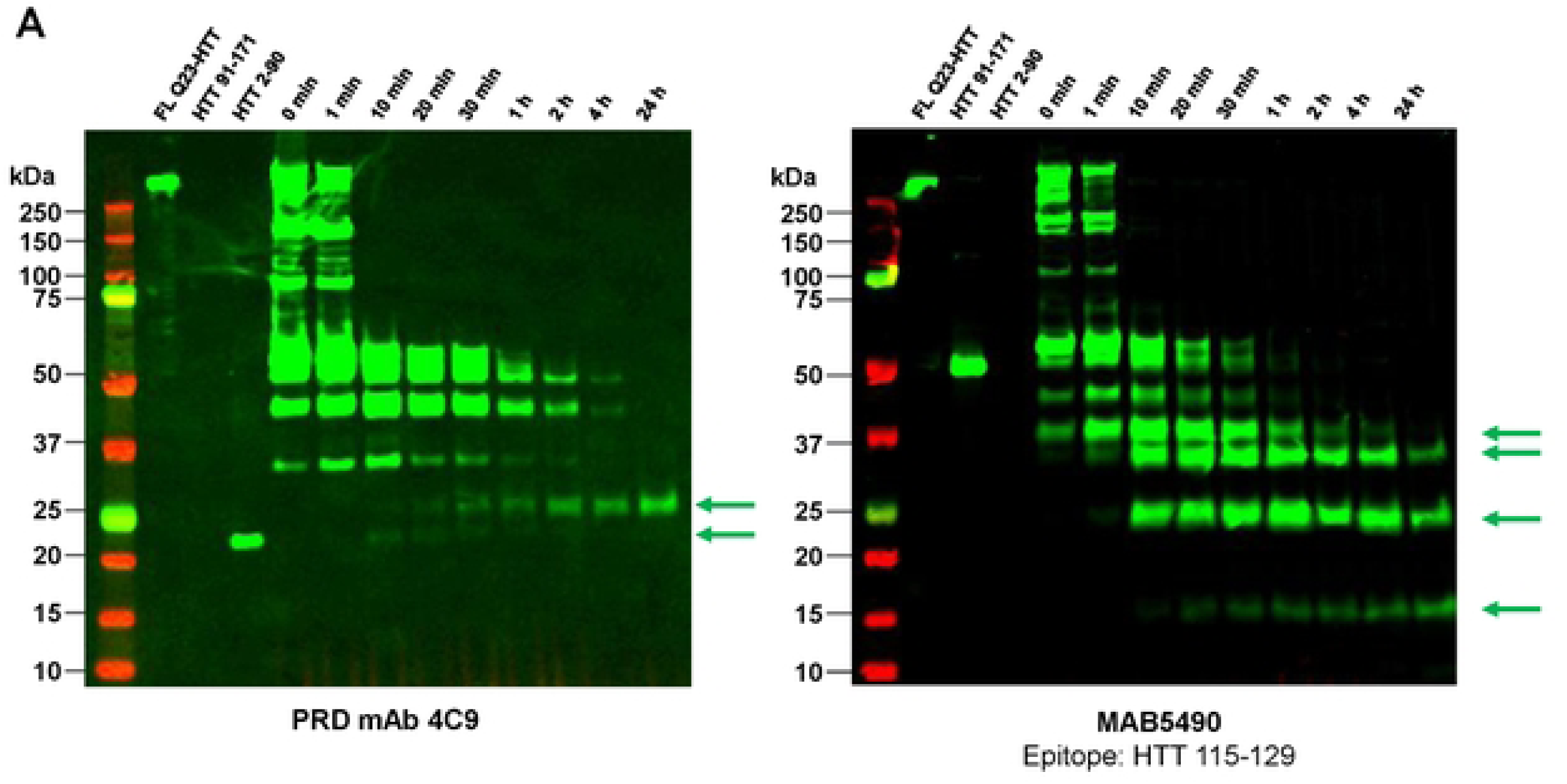
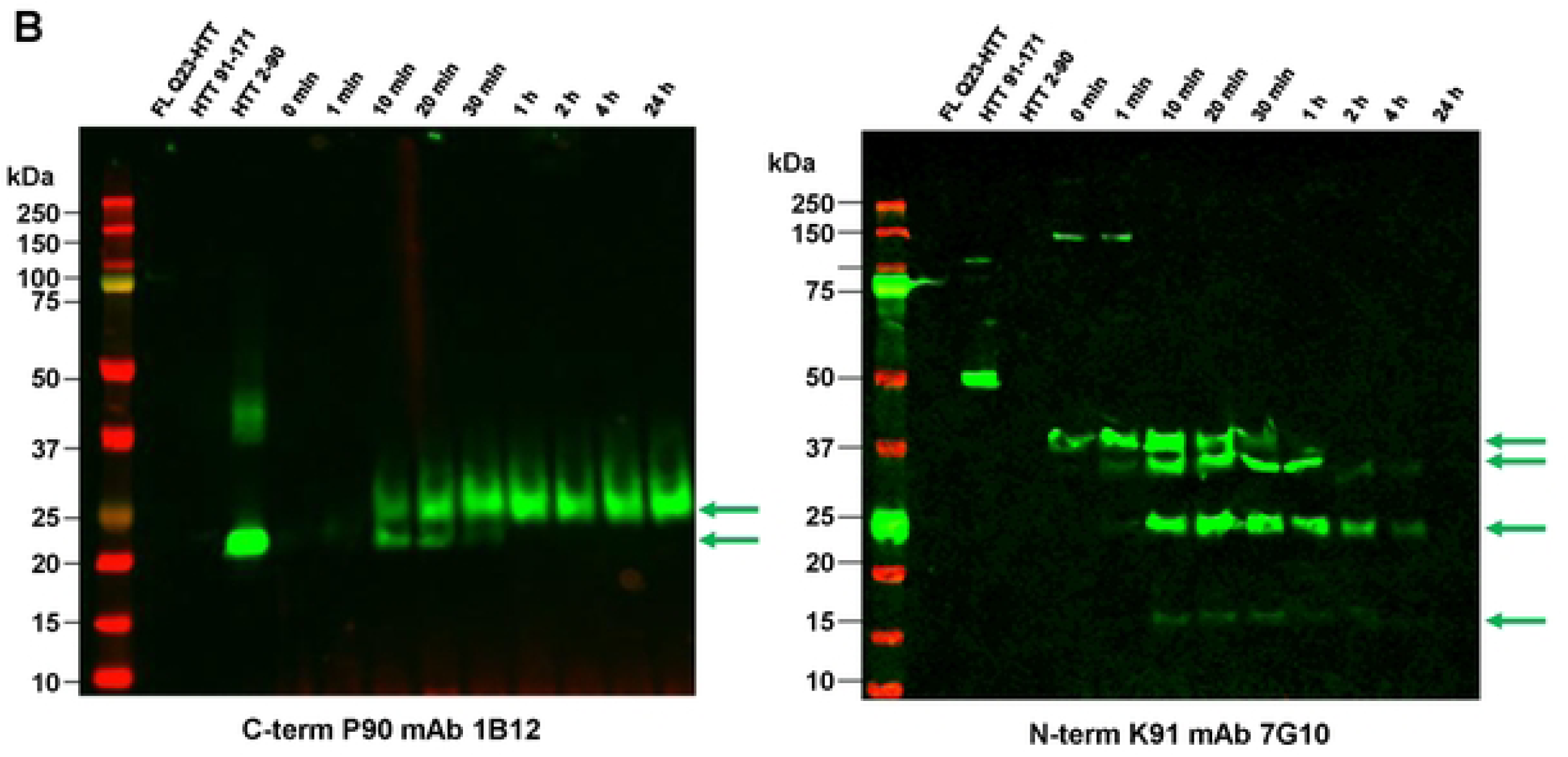
Western blot characterization of Lys-N digest fragments of recombinant FL Q23 HTT. (A) mAb 4C9 and MAB5490; (B) C-term P90 mAb 1B12 and N-term K91 mAb 7G10. Lanes: (1) protein standard ladder; (2) FL Q23 HTT; (3) HTT 1-171 C-term MBP; (4) HTT Q23 Exon1 (aa 2-90); (5-13) crude product of Lys-N digest time course 0-24 hr. Green arrows indicate mAb positive N-term fragments.

The Lys-N digest was compared to Arg-C under comparable conditions (Fig 14). Again, the time-course study revealed a complex fragment profile based on mAb 4C9 (PRD) and MAB5490 (Exon2) characterization. However, the digest did not reveal any 1B12- or 7G10-positive bands, even under increased exposure. This result is consistent with the expectation that Arg-C cleavage does not reveal the C-terminal P90 or N-terminal K91 neoepitope.

**Fig 14.**
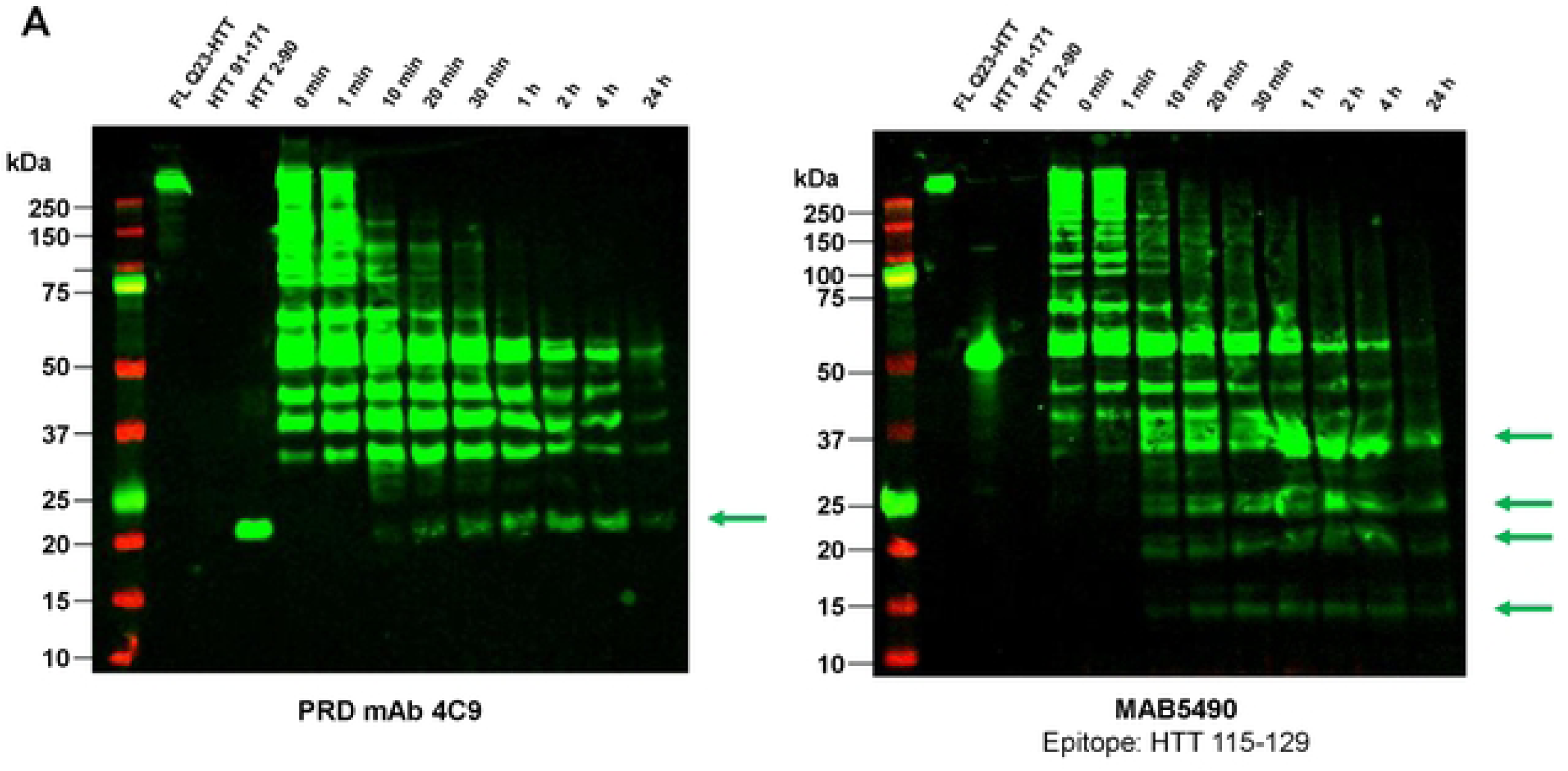
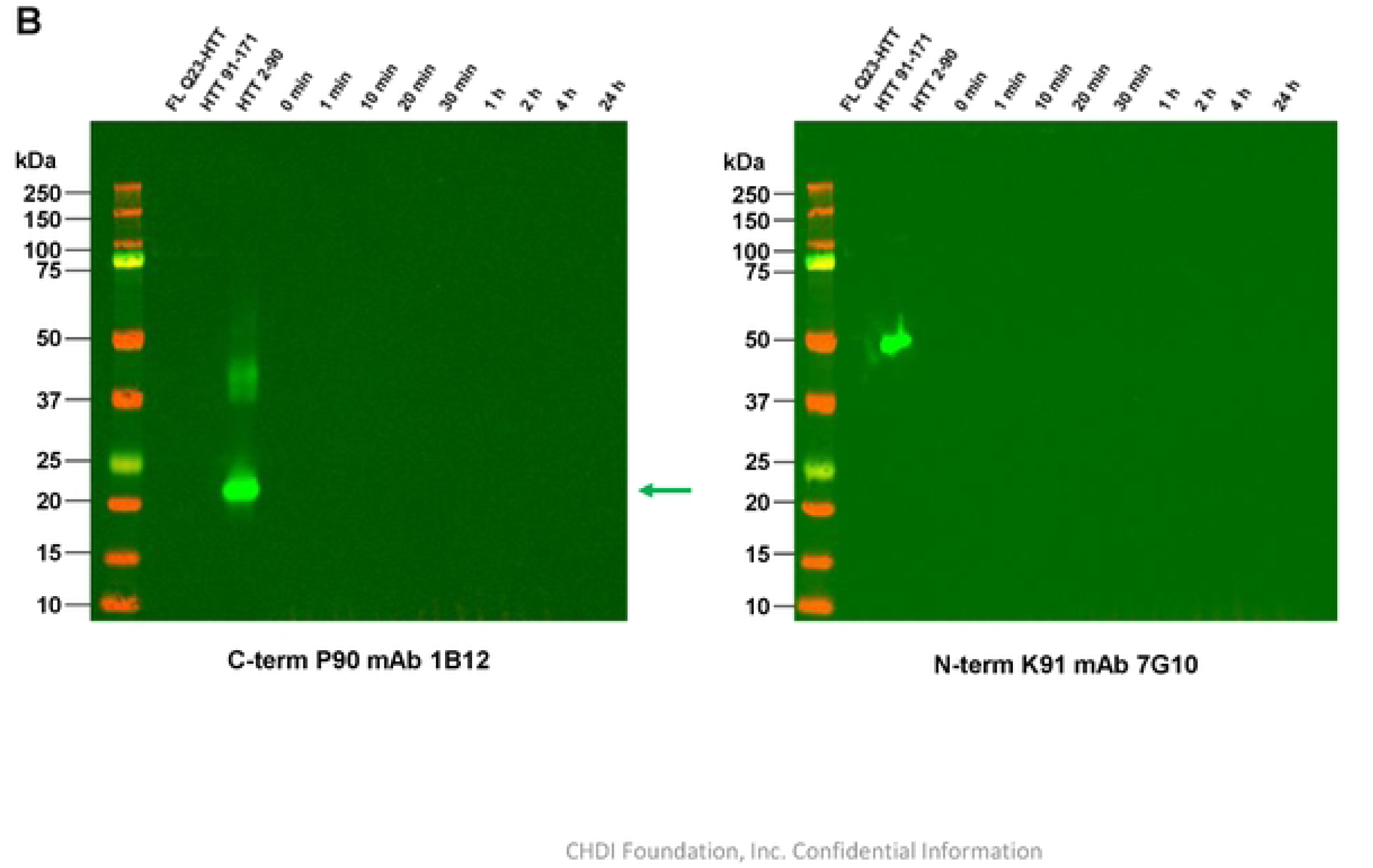
Western blot characterization of Arg-C digest fragments of recombinant FL Q23 HTT. (A) mAb 4C9 and MAB5490; (B) C-term P90 mAb 1B12 and N-term K91 mAb 7G10. Lanes: (1) protein standard ladder; (2) FL Q23 HTT; (3) HTT 1-171 C-term MBP; (4) HTT Q23 Exon1 (aa 2-90); (5-13) crude product of Arg-C digest time course 0-24 hr. Green arrows indicate mAb positive N-term fragments. Exposure time for clone 1B12 and 7G10 was increased and did not reveal mAb positive fragments.

To further validate mAb specificity, Lys-N and Arg-C proteolysis was repeated on human FL Q48 HTT, and the polyQ-containing fragments were isolated by anti-polyQ mAb 3B5H10 pull-down. After recovery of the polyQ-rich N-terminal fragments from the 3B5H10-coated resin, intact MS was performed to confirm that the predicted fragments (Fig 15) were generated: K15–P90 for Lys-N and A2–R89 for Arg-C.

**Fig 15.**
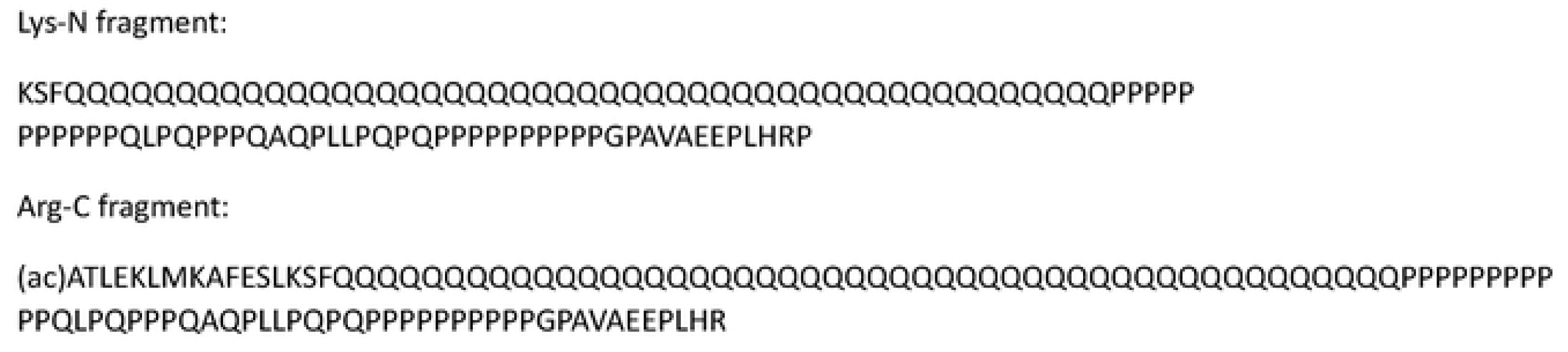
Predicted fragments from pull down with 3B5H10 after digest of FL Q48 HTT with Lys-N or Arg-C.

After Lys-N digestion, an intact mass of 11683.49 Da (theoretical: 11683.61 Da) was observed (Fig 16A), corresponding to the predicted Q48 Exon1 fragment K15–P90. The Arg-C digest produced a Q48 Exon1 fragment with an observed intact mass of 13091.71 Da (theoretical: 13091.28 Da), corresponding to the N-terminally acetylated fragment A2–R89 (Fig 16B). This observed fragment is consistent with the findings of Huang et al. [24], who showed that HTT produced in HEK293 cells is N-terminally processed to yield an acetylated N-terminus. By western blot, the isolated Lys-N and Arg-C HTT fragments both showed the expected reactivity to mAb 4C9, confirming the presence of the PRD domain (Fig 16C). The Lys-N fragment was reactive to P90 mAb 11G2, whereas the Arg-C fragment was not.

**Fig 16.**
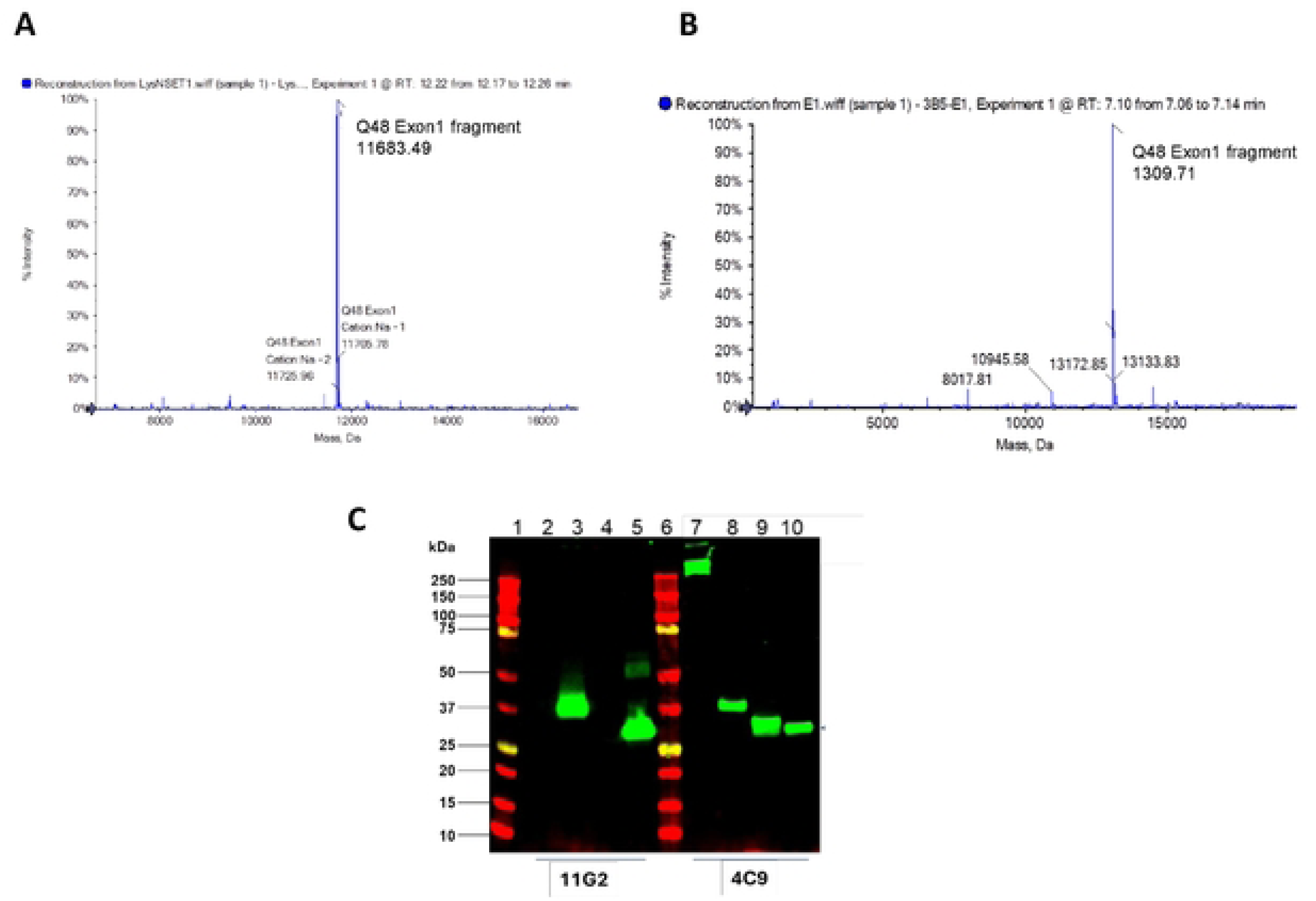
Intact Mass Analysis of 3B5H10 pull-down fragments from FL Q48 HTT after (A) Lys-N digest and (B) Arg-C digest. Western blot characterization of pull-down fragments with mAbs 4C9 and 11G2 (C). Lanes: (1) protein standard ladder; (2) FL Q48 HTT; (3) Lys-N fragment; (4) Arg-C fragment; (5) HTT Q43 Exon1; (6) protein standard ladder; (7) FL Q48 HTT; (8) Lys-N fragment; (9) Arg-C fragment; (10) HTT Q43 Exon1.

## Discussion

In this study, we generated novel recombinant monoclonal antibodies recognizing the HTTexon1 C-terminal P90 neoepitope and the HTTexon2 N-terminal K91 neoepitope—the putative reciprocal cleavage products at the P90–K91 site. We also discovered high-affinity PRD-directed mAbs cross-reactive between mouse and human FL HTT, which will facilitate direct comparison between preclinical mouse models and human biosamples in translational studies.

The P90-specific clones 1B12 and 11G2 demonstrated robust fragment-length selectivity across multiple complementary assay formats. By SPR, both clones showed high affinity for recombinant Exon1 protein and pre-formed HTTexon1 fibrils, with no detectable binding to FL HTT. Western blotting and filter-trap immunoblotting against a panel of HTT fragments terminating at residues from L87 to K99 confirmed that 1B12 and 11G2 selectively detect the P90 C-terminus, with only marginal reactivity toward the adjacent 1–91 fragment observable under sensitive filter-trap conditions. This high selectivity makes these clones particularly well suited for immunoassay-based quantification of HTTexon1 in patient biosamples, an application where distinguishing specific fragment termini from background or larger HTT proteoforms is critical.

We orthogonally validated neoepitope specificity through a proteolysis-mass spectrometry strategy, providing independent, sequence-level confirmation beyond antibody-based methods alone. Lys-N digestion of FL HTT generated the predicted K15–P90 fragment, confirmed by intact mass spectrometry, and this fragment was reactive with the P90 mAb. Conversely, Arg-C digestion, which does not expose the P90 or K91 neoepitopes, produced no reactivity with either neoepitope-directed mAb, providing strong orthogonal evidence for cleavage-site specificity.

Comparative ICC analysis revealed that the P90 clones 1B12 and 11G2 efficiently detect both diffuse monomeric and aggregated HTT 1–90 species with low background, co-localizing with MW8-positive aggregates; this suggests that the P90 neoepitope remains accessible in the aggregated state, a finding that may be relevant for neuropathological applications in HD brain tissue. The P90 rabbit mAbs may therefore be used in combination with the mouse mAb MW8, offering complementary orthogonal detection of the C-terminal P90 neoepitope in both diffuse and inclusion-bearing cellular contexts.

Discrepancies between our MW8 aggregate detection results and those previously reported by Landles et al. prompted a direct, side-by-side comparison of MW8 sourced from CHDI (CHDI-90000942) and from Merck (MABN2529). Using the same panel of HTT fragment constructs, Landles confirmed that Merck-sourced MW8 retains higher C-terminal P90 selectivity, consistent with their earlier published data [4], whereas the CHDI-sourced lot used in our study detected aggregates formed by fragments extending beyond P90 (K91, E93, S95, and K99) (Fig 9). This finding demonstrates that the CHDI and Merck MW8 lots, despite sharing a common hybridoma origin, have measurably drifted from one another. Importantly, both sources remain in active circulation and use within the HD research community. Investigators should therefore be aware that results obtained with MW8 may be source- and lot-dependent, and that findings generated with one source may not be directly reproducible with the other. By contrast, the recombinant P90 mAbs described here (1B12, 11G2) are expressed from a defined, sequenced clone, and their performance is not expected to be subject to this type of source-dependent drift—reinforcing recombinant production as a path to long-term reagent consistency across laboratories.

While the C-terminal P90 neoepitope was robustly detected across all formats evaluated, the biological relevance of the K91 N-terminal neoepitope warrants further investigation. It remains to be established whether endogenous proteolytic cleavage at P90–K91 occurs in human or mouse tissues, and whether this event contributes to normal HTT processing or HD pathology. The availability of the validated 7G10 mAb provides a specific reagent to address this open question in future studies using patient-derived material and knock-in mouse models.

Results from WB, ICC, IHC, MSD and HTRF detection assays employing the P90 neoepitope mAbs 1B12 and 11G2 have been recently reported [25–29], and the availability of these mAbs ensures broad community access to support reproducible, comparable measurements of HTT proteoforms across laboratories. Collectively, the antibodies described here—1B12, 11G2, 27F5, 31C10, and 7G10—address a longstanding gap in the HD field by providing well-characterized, recombinant reagents with defined specificity profiles across multiple detection modalities.

The mAbs are available in the Coriell Institute HD Community Biorepository: anti-HTT neo P90, clone 1B12 (CHDI-90004290); anti-HTT neo P90, clone 11G2 (CHDI-90004291); anti-HTT neo K91, clone 7G10 (CHDI-90004394); anti-HTT, human/mouse PRD cross-reactive, clone 31C10 (CHDI-90004392); anti-HTT, human/mouse PRD cross-reactive, clone 27F5 (CHDI-90004387). The full sequences for the antibodies are provided in the Supporting Information (S4).

## Acknowledgements

The authors thank Jonathan Bard, Deanna Marchionini, and Simon Noble for their editorial input. We also thank Douglas Macdonald for helpful discussions.

---

**Fig S3.**
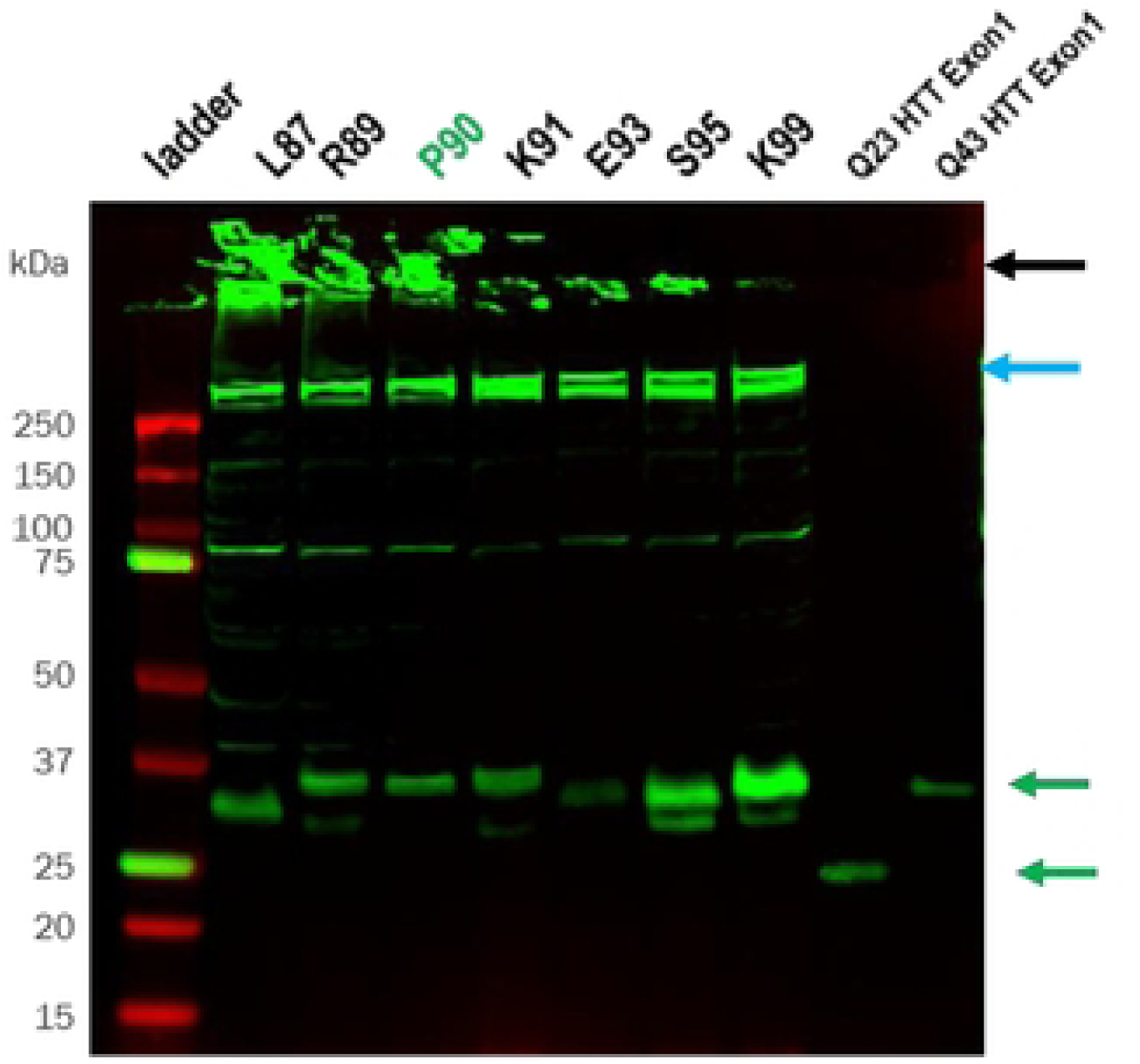
Over expression studies in HEK293. Fig 7 blots re probed with the N17-directed mAb 287 to confirm the identity and expression of the HTT fragments. Lanes: (1) ladder; (2) C-term L87; (3) C-term R89; (4) C-term P90; (5) C-term K91; (6) C-term E93; (7) C-term S95; (8) C-term K99; (9) recombinant HTT Q23 Exonl; (10) recombinant HTT Q43 Exonl. 8lack arrow indicates aggregates. Blue arrow indicates endogenous FL HTT. Green arrows indicate Exonl fragments.

